# Cortical excitatory and inhibitory neuron deficits may underlie the cognitive and social impairments in a mouse model of schizophrenia with exonic *Reln* deletion

**DOI:** 10.1101/2025.07.21.666057

**Authors:** Youyun Zhu, Kanako Kitagawa, Daisuke Mori, Tetsuo Matsuzaki, Taku Nagai, Toshitaka Nabeshima, Sayaka Takemoto-Kimura, Hiroaki Ikesue, Norio Ozaki, Hiroyuki Mizoguchi, Kiyofumi Yamada

## Abstract

Reelin is an essential extracellular matrix glycoprotein involved in the formation of cortical layers and has been associated with several neuropsychiatric conditions, such as schizophrenia (SCZ). To explore its role in brain function and its potential involvement in SCZ, we developed a *Reln* heterozygous deletion (*Reln^del/+^*) mouse model that mimics a genetic deletion observed in a Japanese patient with SCZ. In previous studies, we demonstrated that *Reln^del/+^* mice exhibit cognitive impairments in a visual discrimination test. Here, we found that *Reln^del/+^* mice displayed impairments in social novelty recognition, while social preference remained intact. Immunohistochemical analyses revealed a significant decrease in the numbers of calcium/calmodulin-dependent protein kinase II (CaMKII)-positive glutamatergic pyramidal neurons, gamma-aminobutyric acid (GABA)-ergic interneurons, and parvalbumin (PV)-positive interneurons in the medial prefrontal cortex (mPFC) of *Reln^del/+^* mice. Furthermore, *Reln^del/+^* mice exhibited significant deficits in excitatory spine density and morphology, as well as a decrease number of PV boutons in the mPFC compared to wild-type (WT) mice. Finally, we demonstrated that injection of AAV-R36-Myc virus into the mPFC can improve social novelty impairments in *Reln^del/+^* mice, but no effects on WT control. These findings indicate that *Reln^del/+^* mice could be a valuable model for exploring the neurobiological mechanisms underlying cognitive and social impairments in SCZ. Futhermore, our results with AAV-R36-Myc also suggest the therapeutic potential of Reelin replacement, warranting further investigation as a possible treatment strategy for SCZ.

**Highlights:**

- Reelin deficiency disrupts social novelty, but not social preference, in *Reln^del/+^* mice
- *Reln^del/+^* mice serve as a novel model to investigate impairments of neuronal mechanisms on schizophrenia
- AAV-R36-myc injection rescues social novelty deficits in *Reln^del/+^* mice
- Reelin replacement therapy is a potential treatment for schizophrenia

**Graphical abstract:** 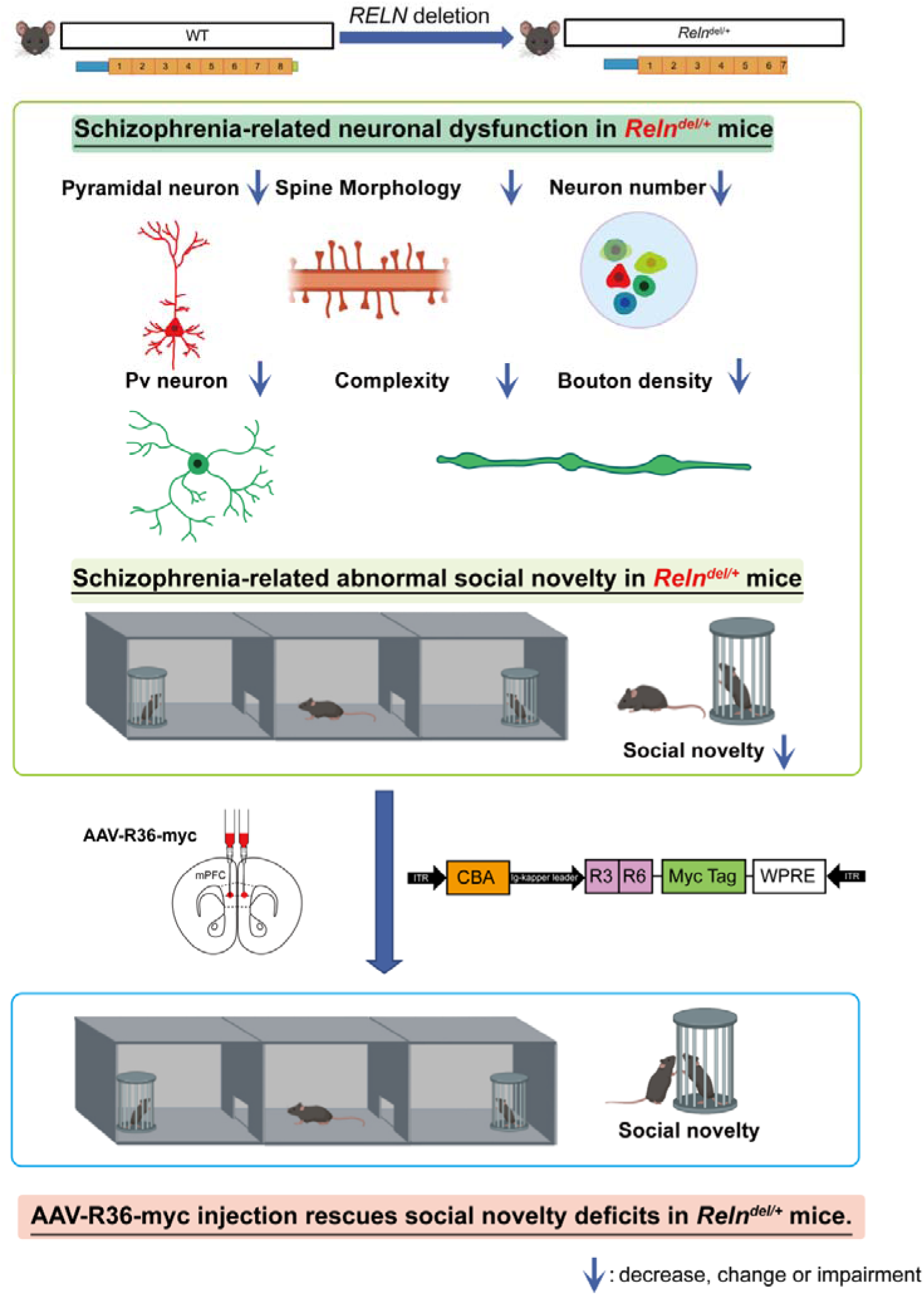

## 1. Introduction

Reelin, an extracellular matrix glycoprotein composed of eight epidermal growth factor (EGF)-like repeat domains secreted by Cajal-Retzius cells, is necessary for cortical layer formation and neuronal migration during brain development [1], [2], [3]. It also promotes the initial growth of axonal and dendrite processes [4]. Reelin is endogenously cleaved at two specific sites—between its second and third, and fifth and sixth EGF-like repeats—resulting in the formation of five fragments [5]. Previous studies have showed that R36 fragment, comprising Reelin’s third to sixth EGF-like repeats of Reelin, which is critical for its signaling and neuronal function, can enhance cognitive function [6],[7],[8]. In the adult brain, Reelin is predominantly produced by gamma-aminobutyric acid (GABA)-ergic interneurons and is a key factor in regulating neuronal plasticity and cognitive function [9], [10], [11].

Reelin signaling enhances GABAergic neurotransmission, promotes synaptic maturation, and upregulates both the expression and activity of α-amino-3-hydroxy-5-methyl-4-isoxazolepropionic acid (AMPA) and N-methyl-d-aspartate (NMDA) receptor subunits [12], [13].

Reelin signaling is dysregulated in several neurological disorders, including schizophrenia (SCZ), autism, depression, and Alzheimer’s disease [14]. In previous studies, it has been reported that patients with SCZ have decreased Reelin and mRNA levels [15].

Reduced Reelin expression in SCZ is associated with structural abnormalities [16], deficits in neural plasticity [10], and impaired gamma oscillation in the human dorsolateral prefrontal cortex (DLPFC) [17].

Previously, we have established a *Reln* deletion mouse model on the C57BL/6J background, modeling a genetic deletion identified in a Japanese patient with SCZ [18]. This deletion corresponds to the one identified in the human patient, allowing us to investigate the effects of reduced Reelin expression [19].

Despite a significant reduction in Reelin mRNA level, heterozygous *Reln* deletion (*Reln^del/+^*) mice did not show any apparent brain malformation [19]. However, they showed a loss of PSD95 puncta in primary cortical neuron cultures, indicating potential synaptic deficits [20]. These synaptic alterations may contribute to brain dysfunctions in this SCZ mouse model, such as cognitive impairments [21]. However, the details of brain dysfunction in the *Reln^del/+^*mouse remain unclear.

In this study, we first used a three-chamber test to assess whether *Reln^del/+^* mice show negative symptom-like behavioral changes characteristic of SCZ. Neuropathological analysis of the mPFC revealed reduced numbers of CaMKII-positive pyramidal neurons, GABAergic interneurons, and PV-positive interneurons, along with significant morphological abnormalities and decreased spine density in pyramidal and PV-positive interneurons. The VGLUT2/VGAT ratio on their soma remained unchanged. Building upon previous findings that a novel construct containing the R36 region induces receptor dimerization and activates downstream Reelin signaling [6], [7], we investigated the effects of AAV-R36-Myc microinjections as an initial step toward evaluating Reelin replacement therapy for SCZ. Notably, R36 treatment effectively rescued social novelty recognition deficits in *Reln^del/+^*mice, supporting its potential as a therapeutic approach for *RELN*-associated impairments.

## 1. Materials and Methods

### 1.1. Animals

The C57BL/6J mice used in this study were sourced from Japan SLC Inc. (Hamamatsu, Japan). *Reln^del/+^* mice on this background were established as previously reported [19].

Age-matched wild-type (WT) littermates served as controls. PV-IRES-Cre mice (B6;129P2-Pvalb*^tm1(cre)Arbr^*/J; stock 008069) were obtained from The Jackson Laboratory (USA). To target PV-positive interneurons in *Reln^del/+^* mice, we crossed PV-IRES-Cre (PV-Cre) mice with *Reln^del/+^* mice to obtain *Reln^del/+^*; PV-Cre mice. The mice were housed in groups under a 12-h dark/light cycle, with food and water available ad libitum. Both male and female adult mice older than 8 weeks were used in the study. All animal experiments were conducted in strict accordance with the Principles for the Care and Use of Laboratory Animals established by the Japanese Pharmacological Society and National Institutes of Health (NIH) Guide for the Care and Use of Laboratory Animals, following review and approval by the Animal Care and Use Committee of Nagoya University.

### 1.1. Social Preference and novelty test

A three-chamber test was performed to evaluate social deficits [22]. The apparatus (60 × 40 × 22 cm) comprised three connected chambers separated by doorways. Mice were habituated to the test room for at least 1 h. During the initial 10-minute habituation session, mice explored the apparatus freely with two empty metal cages placed in the side chambers. In the pre-test phase, two identical non-social objects (golf ball, O1) were placed inside the metal cages. The social test consisted of two phases (Fig.1A). In the social preference phase, a wooden block (O2) was placed in one side chamber as the non-social stimulus, while the opposite contained a social stimulus (an age- and sex-matched WT mouse-1, M1) enclosed in a metal cage. In the last social novelty phase, the test mouse was presented with the familiar mouse used in the social preference phase (M1) in one chamber and a novel social stimulus (an age- and sex-matched WT mouse-2, M2) in the opposite chamber. Between test phases, mice were returned to their home cages for a 5-minute break, and the chamber was thoroughly cleaned with 75% ethanol. Mouse behavior was recorded and analyzed using EthoVision XT 16 software, which quantified the time spent in proximity to the metal cages (nose-to-cup distance ≤ 3.5 cm). Social preference and novelty index scores were calculated using the following formulas:

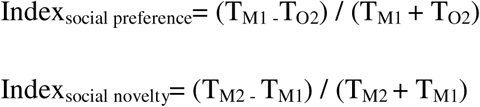

**Fig. 1.**
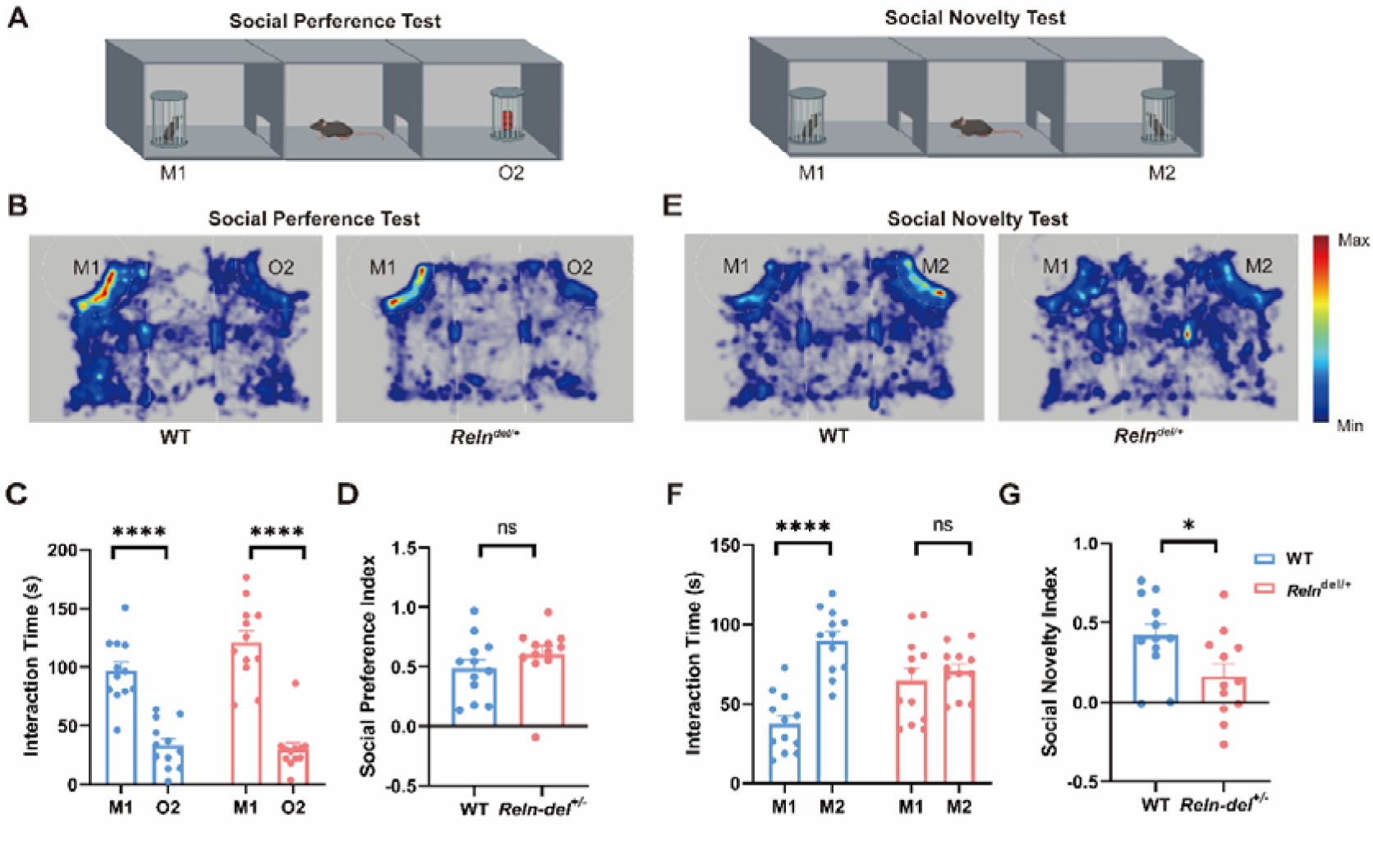
Relndel/+ mice showed impairments in social novelty but in not social preference. A. Schematic of the three-chamber test protocol, including: a 10-min social preference phase with one cage containing a social stimulus (age- and sex-matched WT mouse-1, M1) and the other a non-social object (wooden block, O2); and a 10-min social novelty phase featuring a novel social stimulus (age- and sex-matched WT mouse-2, M2) and a familiar social stimulus (M1). B. Representative heat maps depicting time distribution during the social preference test across mouse groups. C. Interaction time with the social stimulus (M1) versus the non-social stimulus (O2) in WT and *Reln^del/+^* mice. D. The social novelty index comparison between groups. E. Representative heat maps depicting time distribution during the social novelty test across mouse groups. F. Interaction time with novel (M2) and familiar (M1) social stimuli presented in WT vs. *Reln^del/+^* mice. G. The social novelty index comparison between groups. For data represented as mean ± S.E.M (male n = 5-6, female n = 5-6, total n = 10-12 for WT mice, male n = 5-6, female n = 5-6, total n = 12 for *Reln^del/+^* mice); two-way ANOVA For all figures, ns. indicates not significant, *p < 0.05, **p < 0.01, ***p < 0.001, ****p < 0.0001, WT vs. mutant mice.

The difference in time spent between two stimuli (T_O2_ or T_M1_ and T_M1_ or T_M2_) was normalized by their total exploration time.

### 1.2. Plasmid construction

The R36 construct was generated as described in a previous study [6], [7] . In brief, the R36 construct was created by human Reelin repeat regions, R3 (amino acids 1313–1614) and R6 (2339–2618). The resulting R36 construct (VB230918–1815ent, VectorBuilder, Chicago, IL, USA) was inserted into the pTR12.1-MCSW vector, which utilizes a hybrid CMV/chicken β-actin (CBA) promoter and AAV2 terminal repeats to drive expression. For efficient western blot analysis, A Myc-tag was appended to the C-terminus of R36 (Fig.6A).

### 1.3. Virus generation and delivery

HEK293 cells were obtained from ATCC and used between passages 5 and 10. For experiments, HEK293T cells (4 × 10^6^) were seeded in 100-mm dishes (Corning, NY, USA) using Dulbecco’s Modified Eagle Medium (DMEM; Sigma-Aldrich, St. Louis, MO, USA) supplemented with 10% fetal bovine serum (FBS; Gibco, Carlsbad, CA, USA) and 1% Antibiotic-Antimycotic Mixed Stock Solution (100×, stabilized; 09366-44, Nacalai Tesque, Kyoto, Japan) at 37°C with 5% CO_2_. Transfection was performed using AAV plasmids, pHelper and pAAV-DJ (Cell Biolabs, Inc., San Diego, CA, USA) with Opti-MEM medium (Gibco, Carlsbad, CA, USA) and PEI MAX (MW 40,000; Polysciences, Inc., Warrington, PA, USA). The medium containing viral particles was collected after 72 h and centrifuged at 1000 rpm for 5 min at room temperature. The supernatant was discarded and 1 mL of Dulbecco’s phosphate-buffered saline (DPBS; Gibco, Carlsbad, CA, USA) was added to each 1.5 mL tube. After removing the supernatant, 200 μL of DPBS were added. The viral particles were subjected to four cycles of freezing in liquid nitrogen (10 min) and thawing in a 37°C water bath (10 min). Next, 1.5 μL of Benzonase Nuclease HC (71206-25kun, Millipore, Burlington, MA, USA) was added, followed by a 30-minute incubation at 37°C. Samples were centrifuged at 10000 × g for 10 min at room temperature and subsequently filtered with a 0.45 μm filter. The purified virus solution was stored at -80°C.

To analyze spine morphology in cortical pyramidal neurons, a lentivirus encoding enhanced green fluorescent protein (pLLX-EGFP) was used as previously described [20], whereas analysis of PV-positive interneurons was conducted using the AAV-hSyn-FLEX-mCherry virus. AAV-R36-Myc or AAV-Myc was used for the in vivo expression of R36 in the mouse mPFC.

All Mice were anesthetized via intraperitoneal injection of midazolam, medetomidine, and butorphanol mixture (250 mg/kg, intraperitoneally) and then fixed in a stereotaxic frame (David Kopf Instruments, Tujunga, CA, USA). Bilateral injections of pLLX-EGFP (1.7 × 10^8^copies/μL) and pAAV-hSyn-FLEX-mcherry (1.4 × 10^11^ copies/μL) were performed in the mPFC at coordinates: anteroposterior (AP)[=[+1.5mm, mediolateral (ML)[=[±0.5mm, dorsoventral (DV)[=[−2.5mm. An injection of 0.5 μL was administered at each site.

AAV-R36-Myc and AAV-Myc (1 × 10^13^ copies/μL) were microinjected into the mPFC at the same coordinates. A total volume of 5 μL was injected per site. All experiments commenced at least 3 weeks post-injection to ensure the stable viral expression.

### 1.4. Immunohistochemistry and Nissl staining

Immunohistochemistry was performed following the previously [20]. Mice aged 2 – 3 weeks or 8 – 15weeks were anesthetized with isoflurane and perfused intracardially using 4% paraformaldehyde (PFA) in 0.1 M phosphate buffer (PB). Post-fixation was performed in the same solution, followed by cryoprotection sequentially in 20% and 30% sucrose solutions prepared in PB. Coronal brain sections, 20 μm thick, were cut on a cryostat (CM3050S, Leica, Wetzlar, Germany). Sections were then fixed for 5 min in 4% PFA in 0.1 M PB, permeabilized with 0.3% Triton X-100 in PBS for 10 min, and blocked for 1 h in blocking solution (5% serum/PBS with 0.3% Triton X-100). Then the sections were incubated overnight with the primary antibody. Following PBS washes, sections were incubated at room temperature for 1 h with secondary antibody. After additional PBS washes, sections were mounted on adhesive silane (MAS) –coated glass slides (Matsunami, Osaka, Japan) with Fluorescent Mounting Medium (Dako, Santa Clara, CA, USA) and covered with coverslips. Fluorescent images were conducted using confocal laser scanning microscopes (TiE-A1R; Nikon, Tokyo, Japan; and LSM880; Carl Zeiss AG, Oberkochen, Germany). The mPFC were located base on the mouse brain atlas [23].

Nissl staining was performed following protocols [24]. Brain tissues were sectioned at 5 μm thickness on a cryostat, then mounted and observed with a light microscope (FSX100; Olympus, Tokyo, Japan).

### 1.5. Morphological analysis of targeted neurons

Mice aged 2–3 weeks and 7–8 weeks were used in this study. Brain sections were randomly coded during analysis to blind researchers to experimental conditions. Images of neurons in layers 2/3 and 5 of the mPFC, including pyramidal neurons and PV-positive interneurons, were captured using a confocal laser scanning microscope (TiE-A1R; Nikon, Tokyo, Japan) equipped with 100× oil-immersion objective lens. Only well-impregnated neurons that were clearly separated from adjacent labeled neurons were selected for analysis. To quantify spine plasticity or PV bouton numbers, only dendritic or axonal segments longer than 20 μm in the mPFC were analyzed. For pyramidal neurons analysis, spine density was assessed on dendrites located 50–200 µm from the soma (4–5 dendrites per mouse) and expressed as the number of spines per 10 µm dendrite length. Three-dimensional (3D) tracing of all dendrites and spines was performed using the Filament Tracer tool in Imaris (Bitplane, Zurich, Switzerland).

Spine morphology analysis was categorized according to the following criteria:

1. Stubby: spine length < 1 µm;
2. Mushroom: spine length < 3 µm and max_width (head) > mean_width (neck) × 2;

1. Long thin: mean_width (head) ≥ mean_width (neck);
2. Filopodia-like: length >1 µm (no head).

For PV boutons, we focused on en passant boutons (EPB) in the mPFC, which primarily form axospinous synapses and exhibit relative stability in the adult brain [25].

High-magnification images of axonal fragments were analyzed using image analysis software written in MATLAB (EPBscore [25]). Axons from PV-positive interneurons (6–9 neurons per mouse) of *Reln^del/+^*; PV-Cre or PV-Cre control mice in the mPFC were imaged. The number of boutons was automatically quantified using EPBscore software. Additionally, Sholl analysis was performed in ImageJ Fiji to assess the total of dendritic length, the number of dendritic intersections and the number of dendritic branches per neuron (5 neurons per mouse).

The number of VGLUT2- and VGAT-immunoreactive puncta on the soma of PV-positive and pyramidal neurons was quantified, as previously described, and the VGLUT2/VGAT ratio was analyzed using Fiji software [26], [27]. Images were acquired from 10–15 neurons per mouse in the mPFC using a confocal laser scanning microscope (TiE-A1R; Nikon, Tokyo, Japan) equipped with a 100× oil-immersion objective lens. Protein levels were quantified by measuring optical density after background correction. Background subtraction was performed using a rolling ball radius of 50, followed by Gaussian blur filtering (σ = 1). The neuronal outline was expanded by 0.5 μm from the soma surface to establish the region of interest (ROI) between the original and expanded outlines. The ROI was binarized, and puncta were quantified after filtering by particle size (0.1–1 μm²) and circularity (0.3–1) [27].

### 1.6. Western blotting

Western blotting was performed on tissue from 8–12-week-old mice, following previously established protocols [28]. Briefly, mPFC tissue was dissected following anatomical coordinates provided in the Mouse Brain Atlas [23] . To extract proteins, sonicated brain tissues were incubated in lysis buffer containing 1% SDS, complete protease inhibitor cocktail (11873580001, Roche, Switzerland) and phosSTOP phosphatase inhibitors (4906837001, Roche, Switzerland) at 99°C for 3 min. After centrifugation at 15,000 rpm for 20 min, protein concentrations in the supernatant were determined using the DC Protein Assay Kit (5000111JA, Bio-Rad Laboratories, Hercules, CA, USA).

Proteins were denatured by boiling in sample buffer containg Tris–HCl (0.5 M, pH 6.8), SDS (1%), glycerol (30%), dithiothreitol (DTT, 0.93%) and bromophenol blue (0.0012%). The denatured proteins were then separated on a 10% SDS–polyacrylamide gel and transferred onto a polyvinylidene difluoride (PVDF) membrane (IPvH00010, Millipore, Darmstadt, Germany). The membranes were blocked at room temperature for 1 h with Blocking One-P (05999-84, Nacalai Tesque, Kyoto, Japan) and then incubated with the primary antibody at 4°C overnight. After three washes (10 min each) with Tris-buffered saline containing 1% Tween-20, membranes were incubated with horseradish peroxidase (HRP)-conjugated secondary antibodies at room temperature for 1 h. Immune complexes were visualized using ECL Plus Western Blotting Detection Reagent (RPN2236, GE Healthcare, Pittsburgh, PA, USA). Band intensities were quantified using a LuminoGraph I imaging system (Atto Instruments, Tokyo, Japan). Western blot data are expressed as relative fold changes in protein expression compared to the control group. To enhance antibody-antigen binding, primary and secondary antibodies were diluted using Can Get Signal Solutions 1 and 2 (NKB-101, Toyobo, Osaka, Japan), respectively.

### 1.7. Statistical analysis

Results are presented as the mean ± standard error of the mean (SEM). Statistical analyses were conducted using GraphPad Prism8 (GraphPad Software Inc., San Diego, CA, USA). Differences between two groups were evaluated using the student’s t test. Statistical comparisons across groups were conducted using one- or two-way ANOVA, followed by appropriate post hoc tests (Tukey’s, Bonferroni’s, or Sidak’s) for multiple comparisons. No data was excluded from the analysis. All collected data were included and analyzed.

## 2. Result

### 2.1. *Reln^del/+^* mice showed impairments in social novelty but not social preference

We assessed social preference and social novelty in the three-chamber test. To evaluate social preference in *Reln^del/+^* mice, we measured the time spent investigating a wooden block (O2), as defined in the Methods section, compared to a social stimulus mouse (M1) during the social preference test phase. Subsequently, in the social novelty phase, O2 was exchanged for a novel social stimulus mouse (M2), while M1 remained as the familiar social stimulus mouse (Fig. 1A). Representative heatmaps of the mice’s trajectories are shown in Fig.1B and 1E.

*Reln^del/+^* mice showed normal social preference behavior, similar to WT controls (Fig. 1C,1D). When presented with a novel (M2) and familiar (M1) social stimulus, WT mice preferentially interacted with M2 for a longer duration, reflecting intact social novelty recognition. In contract, *Reln^del/+^* mice did not show a preference for M2 (Fig. 1F).

Accordingly, the social novelty index was significantly decreased in *Reln^del/+^* mice relative to WT controls (Fig. 1G). These findings suggest that *RELN* deletion selectively impairs social novelty recognition while maintaining social preference. This behavior phenotype may reflect social cognitive impairments observed in SCZ.

### 2.1. *Reln^del/+^* mice showed decreased numbers of CaMKII-positive pyramidal neurons and GABAergic interneurons in the mPFC

Previous studies have revealed that Reelin signaling significantly contributes to the migration of pyramidal neurons and cortical layer formation during development [29]. At 2–3 weeks of age, no significant differences in body weight or brain weight were observed between WT and *Reln^del/+^* mice. However, *Reln^del/del^* mice were significantly smaller than WT littermates (Fig. S1A–S1D) and showed obvious abnormalities in cortical layer structure (Fig. S1G), which were not apparent in *Reln^del/+^* mice. Notably, *Reln^del/+^* mice exhibited a thinner layer2/3, but not layer 5 compared to the WT controls (Fig. S1E–S1F). Consistent with this, we measured the pyramidal neuron count in the mPFC of *Reln^del/+^* mice. Our results demonstrated that the number of pyramidal neurons in mPFC layer 2/3, but not that in layer 5 in *Reln^del/+^* mice compared to WT controls (Fig. 2). These findings were consistent with the layer structure results (Fig. S1).

**Fig. 2.**
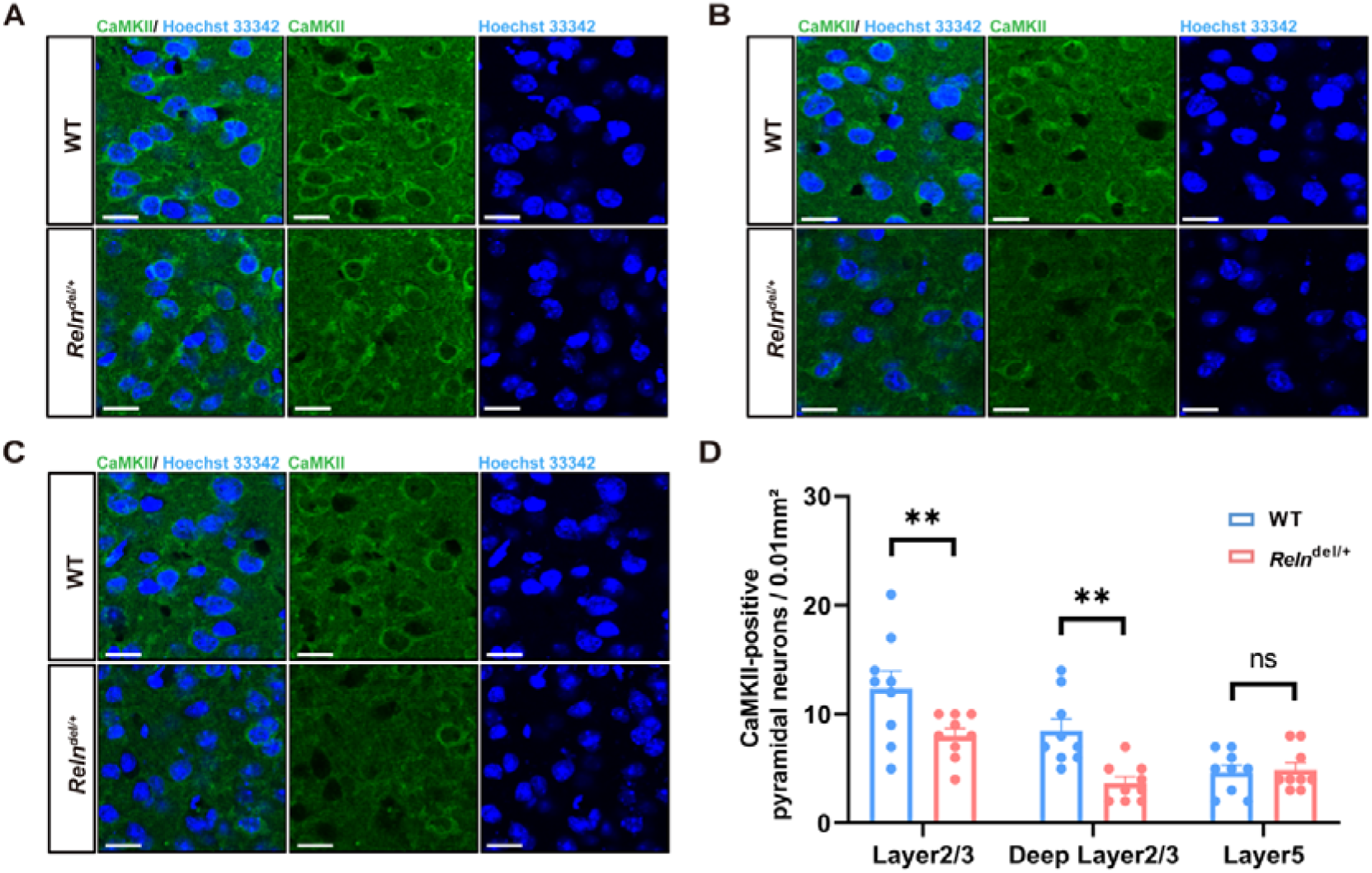
*Reln^del/+^* mouse showed a decreased number of CaMKII-positive pyramidal neurons in the mPFC. A-C. Representative images of CaMKII-positive pyramidal neurons (green) in layers 2/3, deep layer 2/3, and layer 5 of the mPFC from WT and *Reln^del/+^* (9 slices from 3 mice per group). Scale bar, 10μm. D. CaMKII-positive pyramidal neuron numbers per layer in the mPFC. For data represented as mean ± SEM; one-way ANOVA. For all figures, ns.indicates not significant, **p < 0.01, WT vs. mutant mice.

Additionally, the number of GABAergic neurons was significantly decreased in *Reln^del/+^* mice (Fig. 3A, 3B). Since adulthood, Reelin is predominantly produced by GABAergic interneurons [29], we further analyzed the number of Reelin-positive GABAergic interneurons (Reelin-GABA neurons) in the mPFC. A notable reduction was found in Reelin-GABA neurons but not Reelin-negative GABAergic neurons in the mPFC of *Reln^del/+^*mice compared to WT controls (Fig. 3C, 3D). These structural changes in pyramidal and GABAergic neuronal populations may contribute to social behavior deficits observed in *Reln^del/+^* mice, which provide insight into the neurobiological mechanisms underlying SCZ-associated impairments.

**Fig. 3.**
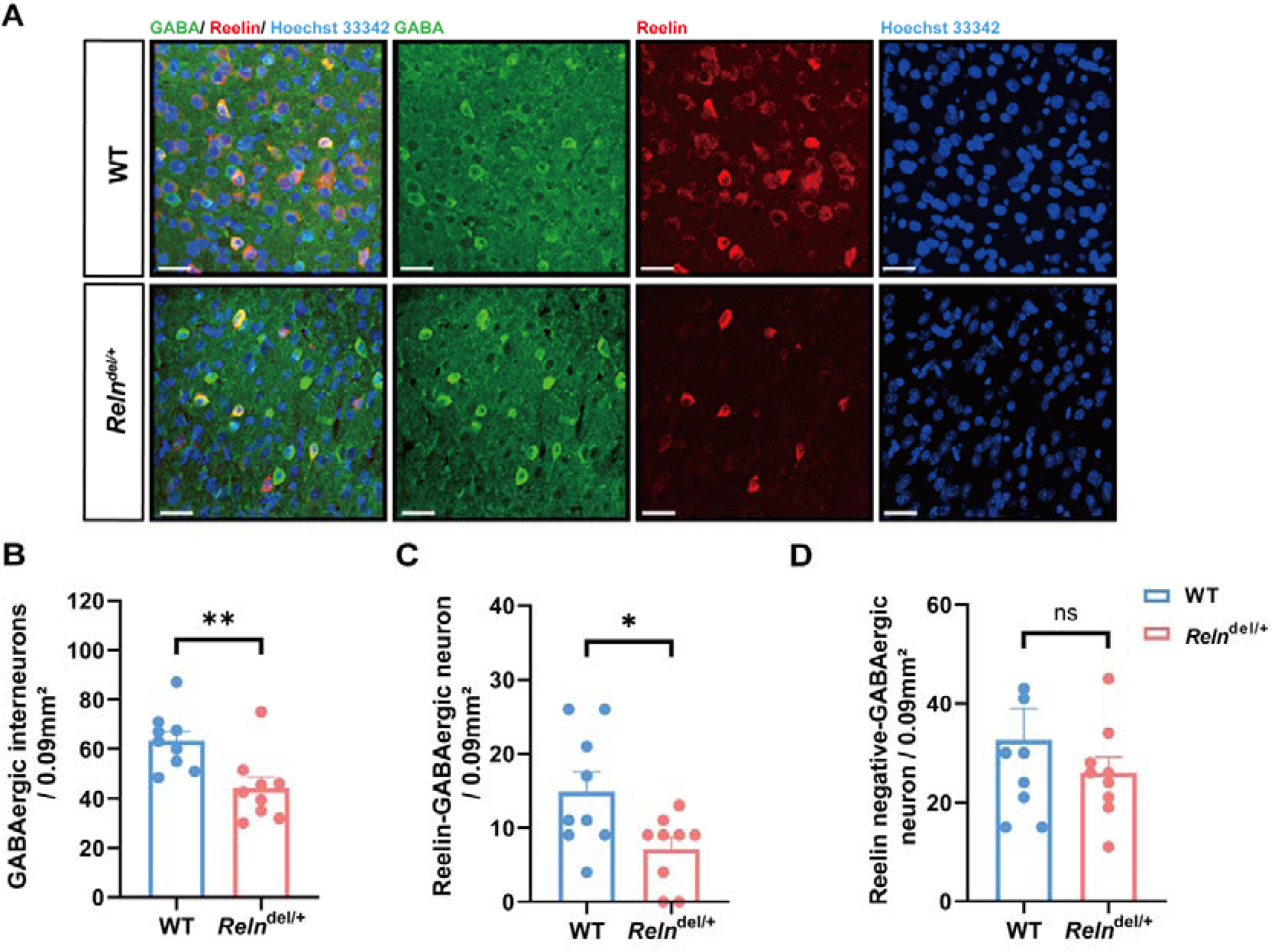
*Reln*^del/+^ mice exhibit reduced GABAergic neuron numbers in the mPFC. A. Representative images showing GABAergic interneurons (green) and Reelin expression in the mPFC of WT and *Reln^del/+^* mice (n=9 slices from 3 mice per group). Scale bar, 30 μm. B. GABAergic interneuron numbers in the mPFC of WT and *Reln^del/+^* mice. C. Reelin-GABAergic neuron numbers in the mPFC of WT and *Reln^del/+^* mice.D. Reelin negative-GABAergic neuron numbers in the mPFC of WT and *Reln^del/+^* mice. For data represented as mean ± SEM; t tests. For all figures, ns.indicates not significant, *p < 0.05, **p < 0.01, WT vs. mutant mice.

### 2.2. *Reln*^del/+^ mice exhibited abnormal spine morphology in deep layers 2/3 and 5 of the mPFC

Pyramidal neurons in the PFC are crucial for the control of social behavior [30], [31], [32], In SCZ, reduced basal dendritic spine density has been detected in deep layer 3 of the DLPFC [17], [33]. Similarly, a well-established Reelin-deficient mouse model, heterozygous reeler mouse (HRM), show reduced spine density on oblique dendrites in layer 5 pyramidal neurons [34]. Based on these findings, we hypothesized that reduced spine density in layers 2/3 and 5 might be associated with the social novelty impairments observed in *Reln^del/+^* mice. To visualize pyramidal neurons, we used a lentiviral construct based on the pLLX vector to express EGFP. The EGFP-labeled neurons demonstrated normal dendritic morphology, with the majority of spines displaying clear head and neck structures (Fig. 4A, 4D). Fluorescent imaging of EGFP-labeled neurons in layers 2/3 and 5 of the mPFC revealed a significant reduction in spine density in *Reln^del/+^* mice compared to WT controls (Fig. 4B, 4E). Across four major dendritic spine subtypes—mushroom, stubby, long thin, and filopodia—*Reln^del/+^*mice consistently exhibited a reduced number of pyramidal neuron spines compared to WT mice (Fig. 4C, 4F). Despite these morphological abnormalities, the excitatory/inhibitory (E/I) balance in pyramidal neurons, as measured by the ratio of VGLUT2 to VGAT (VGLUT2/VGAT ratio), remained unchanged (Fig. S2). These data suggest observed spine abnormalities may lead to reduced neuronal activity without disrupting the local E/I synaptic balance in the mPFC.

**Fig. 4.**
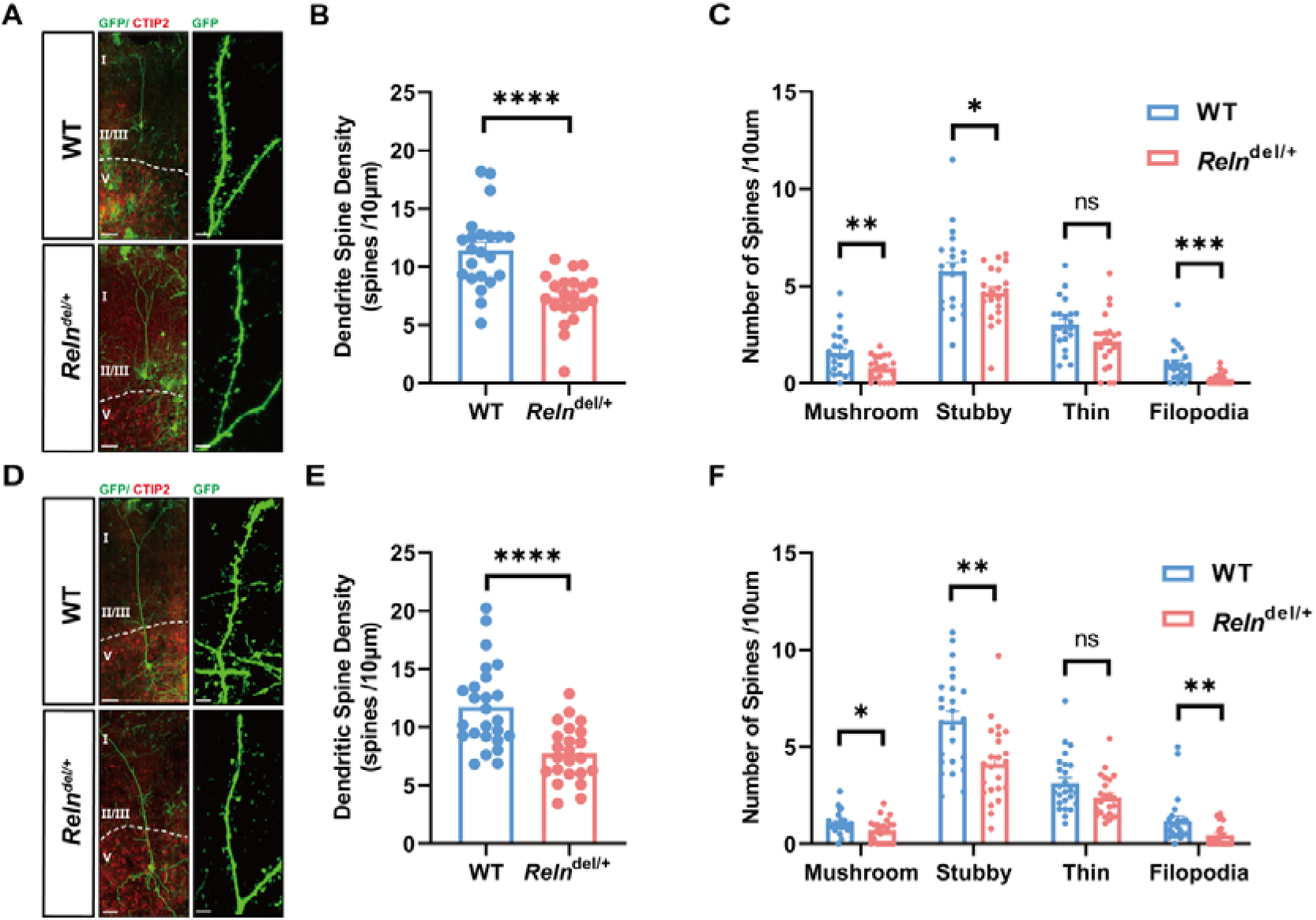
*Reln*^del/+^ mice showed abnormal spine morphology in the deep layer 2/3 and 5 of the mPFC. A. Representative images of GFP-transfected pyramidal neurons in layer 2/3 of the mPFC from WT and *Reln^del/+^* mice (n = 22 neurons from 5 mice per group). A multicolor confocal image taken from a mPFC region that was immunostained against Ctip2 (Red, marker for layer 5), indicating the clear boundaries of the layers. Scale bars, 50 μm (left) and 3 μm (right). B. Dendrite spine density of layer 2/3 pyramidal neurons in the mPFC of WT and *Reln^del/+^* mice. C. Spine morphology analysis in the layer 2/3 of the mPFC of WT and *Reln^del/+^* mice. D. Representative images of GFP-transfected pyramidal neurons in layer 5 of the mPFC from WT and *Reln^del/+^* mice (n = 23–25 neurons from 5 mice per group). A multicolor confocal image taken from a mPFC region that was immunostained against Ctip2 (Red, marker for layer 5), indicating the clear boundaries of the layers. Scale bars, 50 μm (left) or 3 μm (right). E. Dendrite spine density of layer 5 pyramidal neurons in the mPFC of WT and *Reln^del/+^* mice. F. Spine morphology analysis in the layer 5 of the mPFC of WT and *Reln^del/+^* mice. For data represented as mean ± SEM; t tests. For all figures, ns.indicates not significant, *p < 0.05, **p < 0.01, ***p < 0.001, ****p < 0.0001, WT vs. mutant mice.

### 2.3. *Reln^del/+^*; PV-Cre mice showed reduced complexity of PV-positive interneuros in the mPFC

PV-positive interneuros, characterized by fast-spiking activity, are GABAergic neurons that provide inhibitory regulation over cortical and subcortical circuits and are considered critical contributors to the pathophysiology of SCZ [35]. To investigate the effects of *RELN* deletion on PV-positive interneurons, we generated *Reln^del/+^*; PV-Cre mice (Fig. 5A). To selectively visualize PV-positive interneurons and assess their morphology, we administered AAV-hSyn-FLEX-mcherry into the mPFC of *Reln^del/+^*; PV-Cre mice. Our analysis revealed a marked reduction in PV-positive interneuron numbers in the mPFC of *Reln^del/+^*; PV-Cre mice compared to WT; PV-Cre control mice (Fig. 5B, 5C). To examine PV-positive interneurons complexity, we performed Sholl analysis as previously described [28]. This analysis revealed a significant decrease in total dendritic length (Fig. 5E), dendritic branches (Fig. 5F) and dendritic intersections (Fig. 5G) in *Reln^del/+^*; PV-Cre mice relative to WT; PV-Cre controls. These findings indicate that complexity of PV-positive interneurons is markedly reduced in the mPFC of *Reln^del/+^* mice.

**Fig. 5.**
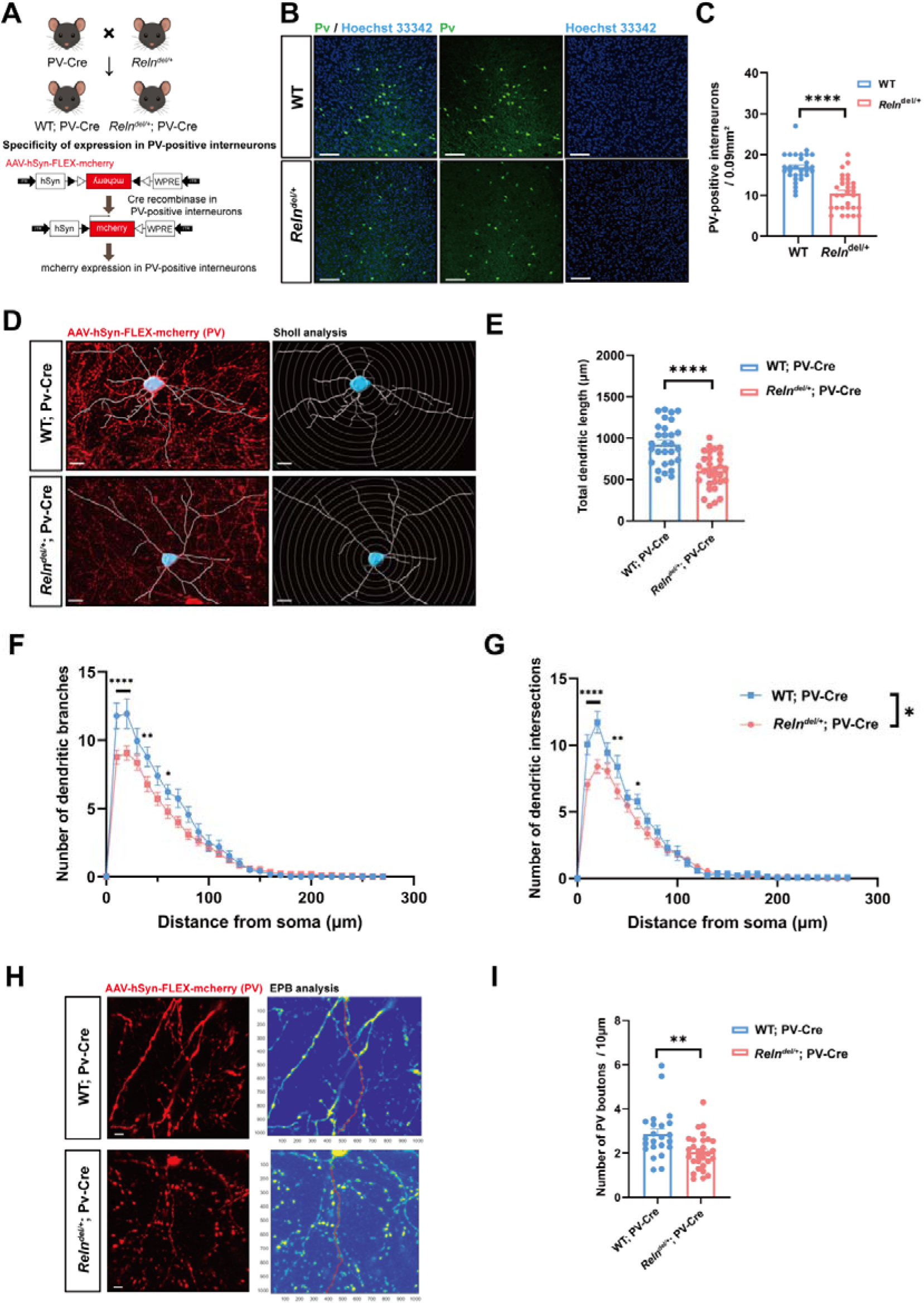
*Relndel/*+; PV-Cre mouse showed a decreased complexity, neuron and bouton numbers in PV-positive interneurons of mPFC. A. Schematic showing the transgene construction of *Reln^del/+^*; PV-Cre transgenic mice. *Reln^del/+^*and PV-Cre mice were crossed to generate *Reln^del/+^*; PV-Cre mice. B. Representative images of PV-positive interneurons (green) in the mPFC of WT and *Reln^del/+^* mice (n= 27-32 slices from 3-4 mice per group) . Scale bar, 100 μm. B. The numbers of PV-positive interneurons in the mPFC of WT and *Reln^del/+^* mice. C. Representative images of PV-positive interneurons (red) analyzed by Sholl analysis in the mPFC of WT and *Reln^del/+^* mice (n=15 slices from 3 mice per group). Scale bar, 10μm D. The total dendritic length of PV-positive interneurons in the mPFC of WT and *Reln^del/+^* mice (sum of dendritic distances from the end of dendric to soma). E. The numbers of dendritic branches of PV-positive interneurons in the mPFC of WT and *Reln^del/+^* mice. F. The numbers of dendritic intersections of PV-positive interneurons in the mPFC of WT and *Reln^del/+^* mice. G. Representative images of PV boutons (red) in the mPFC of WT and *Reln^del/+^* mice (n= 23-29 slices from 3-4 mice per group). Scale bar, 3 μm. H. PV boutons numbers in the mPFC of WT and *Reln^del/+^* mice. For data represented as mean ± SEM; two-way ANOVA and t tests. For all figures, *p < 0.05, **p < 0.01, ****p < 0.0001, WT vs. mutant mice.

Beyond these morphological changes, we also detected a decrease in the number of PV boutons in *Reln^del/+^*; PV-Cre mice. These boutons, which represent a major class of cortical presynaptic terminals, typically appear as swellings along the axonal shaft [36] (Fig. 5H, 5I). Despite these morphologic alterations, the E/I balance, measured by the VGLUT2/VGAT ratio on PV-positive interneurons’ soma, remained unchanged compared to WT mice (Fig. S2).

### 2.4. Reelin fragment R36 rescued social novelty deficits in *Reln^del/+^* mice

Given the observed structural and social behavioral deficits in *Reln^del/+^* mice, we next examined whether restoring Reelin could reverse these impairments. To this end, we tested the effect of R36, a Reelin fragment containing the third and sixth EGF-like repeats, on social behavior. To determine whether R36 (Fig. 6A) can improve the social deficits observed in *Reln^del/+^* mice, we injected either AAV-R36-Myc or AAV-Myc (mock, control) into the mPFC. Western blotting confirmed robust viral expression (Fig. 6B). In the social novelty test, both *Reln^del/+^* mice and WT mice treated with AAV-R36-Myc showed normal social preference, similar to control mice (Fig. 6D, 6E). We confirmed R36 effectively rescues social novelty deficits, since *Reln^del/+^* mice treated with AAV-R36-Myc exhibited a marked preference for the novel mouse, whereas *Reln^del/+^* mice treated with AAV-mock-Myc did not (Fig. 6F, 6G). These findings suggest that Reelin replacement with R36 effectively restores social novelty recognition in *Reln^del/+^*mice, highlighting its potential as a therapeutic strategy for diseases of *RELN*-associated deficits.

**Fig. 6.**
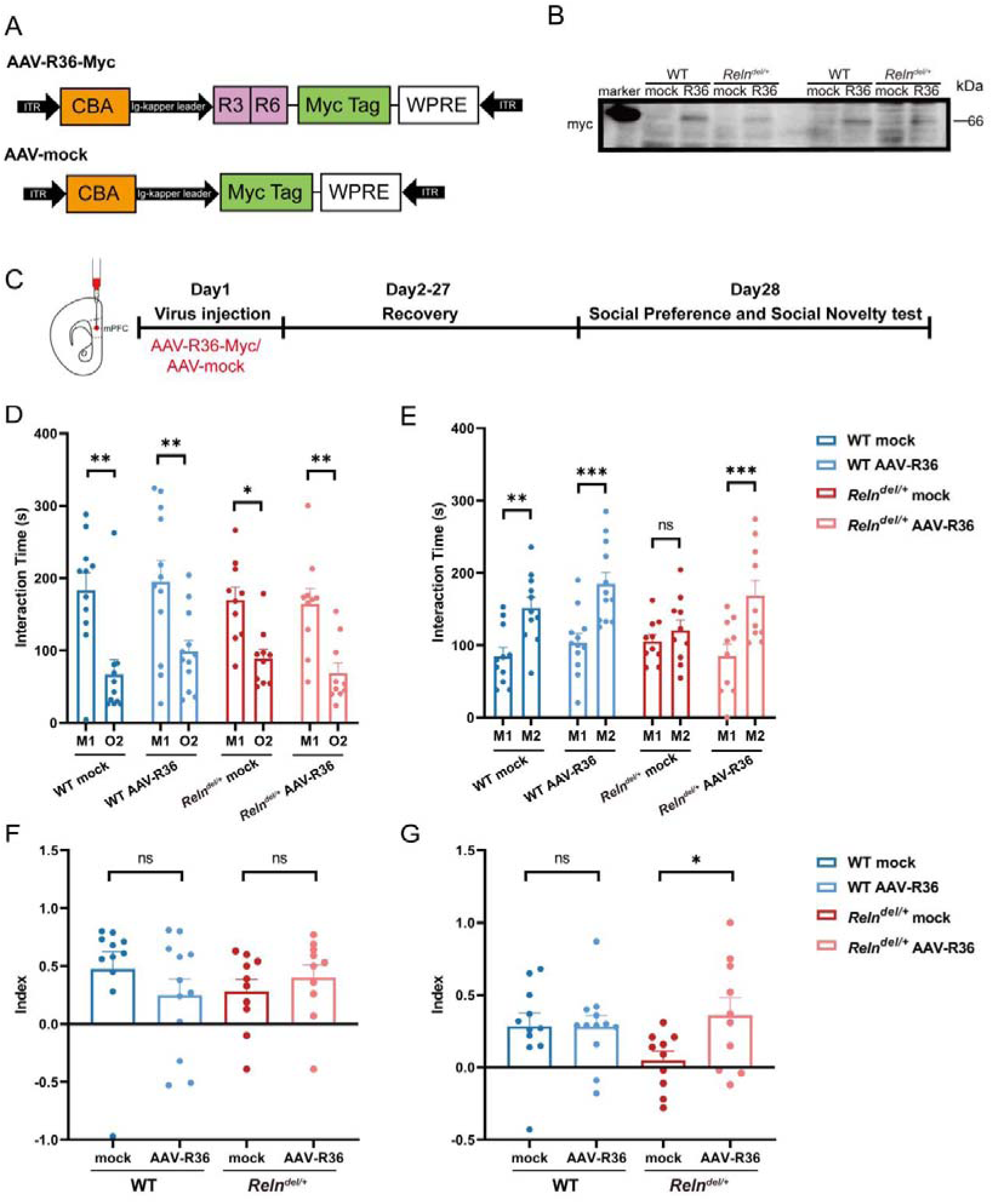
Reelin fragment R36 can rescue social novelty abnormality in *Reln^del/+^* mice. A. Schematic of AAV-R36-Myc and AAV-mock-Myc constructs. The R36 fragment was cloned into the pTR12.1-MCSW vector, with a C-terminal Myc-tag for detection by western blot. B. Western blot showing Myc-tag expression in the mPFC of WT and *Reln^del/+^* mice injected with AAV-R36-Myc or AAV-mock-Myc. R36 expression (∼66 kDa) was detected only in AAV-R36-Myc-injected mice. C. Experimental timeline and design. D. Time spent interacting with social (M1) versus non-social stimuli (O2) during social preference test in WT and mice *Reln^del/+^* mice with AAV-R36 or mock treatment. E. The social preference index comparison between groups. F. The time spent interacting with novel (M2) and familiar (M1) social stimuli during social novelty test in WT and mice *Reln^del/+^* mice with AAV-R36 or mock treatment. G. The social novelty index comparison between groups. For data represented as mean ± S.E.M (male n = 5-6, female n = 5-6, total n=10-12 for WT mock or AAV-R36 mice; male n = 5-6, female n = 5-6, total n=10-12 for *Reln^del/+^* mock or AAV-R36 mice); two-way ANOVA, ordinary one-way ANOVA and t tests. For all figures, ns. indicates not significant, *p < 0.05, **p < 0.01, ***p < 0.001, mock vs. AAV-R36 mice.

## 3. Discussion

Despite recent progress in elucidating the genetics of SCZ, the underlying neurobiological substrates and neural circuits remain largely unknown. The *RELN* gene has become the focus of interest, with accumulating evidences suggesting it as the causative gene of the major neurological symptoms in the SCZ [25], [37], [38]. In this study, we used *Reln^del/+^* mice, a novel *Reln* deletion mouse model based on a Japanese SCZ patient with a specific *RELN* exonic deletion [19], to explore how reduced Reelin levels affect mPFC function and social behavior.

Reduced levels of Reelin have been associated with increased vulnerability to neuropsychiatric conditions, as reported in previous studies [1], [2], [14]. Compared to other Reelin-deficient mouse models, the *Reln^del/+^* mice better represent the Japanese SCZ patient carrying the *Reln* deletion. Furthermore, unlike other Reelin-deficient mice such as the Orleans reeler mice of the BALB/c strain, our *Reln^del/+^* mouse model was generated using the C57BL/6J background, which is commonly utilized in SCZ behavioral research. Notably, Reelin expression in the mPFC of *Reln^del/+^*mice was about 60% lower than that in WT mice (Fig. S3), which suggests a significant reduction in Reelin levels specifically in the mPFC, a critical area involved in social and cognitive functions [39].

To assess social behavior, we used a modified three-chamber test including both social preference and social novelty phases [22]. In the social preference phase, *Reln^del/+^* mice behaved similarly to WT mice, spending more time with a social stimulus mouse than with a novel object. During the social novelty phase, *Reln^del/+^* mice failed to discriminate between familiar and novel social stimuli, implying a specific impairment in social recognition potentially linked to the novel *RELN* mutation.

Reelin regulates dendrite development in the brain [10], and spine density reductions have been reported in Reeler mice and in SCZ patients [32], [40]. In particular, layer 2/3 and 5 pyramidal neurons are critical for brain interregional communication and social information processing [33]. The basal dendrites of layer 2/3 project within or across hemispheres, while layer 5 oblique dendrites receive projections from other regions [32]. Our findings showed layer-specific reductions in pyramidal neuron number and spine density in of *Reln^del/+^*mice, consistent with observations in SCZ patients and Reeler mouse model [33], [41]. These changes may reflect disrupted Reelin signaling, which regulates spine actin dynamics [42]. Notably, mature spines (mushroom-type and stubby-type), which are crucial for stable excitatory synapses [43], were also reduced. These findings are linked to behavioral changes like learning and memory and might reflect the underlying structural changes enabling neural plasticity [44].

Interestingly, while spine density was reduced in both layers 2/3 and 5, the number of CaMKII-positive pyramidal neurons decreased only in layers 2/3 but not layer 5. The selective reduction of pyramidal neurons in layers 2/3 may be because these neurons are more sensitive to Reelin signaling. Alternatively, it may be associated with the difference of wiring to other brain areas as mentioned above. Reelin is essential for cortical development, and its deficiency leads to disrupted layering, inversion, and potential neuronal loss. [45]. Pyramidal neurons in the mPFC exhibit rapid dendritic growth and spine formation during the first postnatal month.[46]. In our study, we observed clear abnormalities in cortical layer formation in both *Reln^del/+^* mice and *Reln^del/del^* mice at 2-3 weeks (Fig. S1). Our findings indicate the importance of Reelin signaling in the early formation of cortical layers, which may be essential for development and functional integration of pyramidal neurons in adulthood. The disruption of cortical structure during this sensitive period may underlie the reduced number of CaMKII-positive pyramidal neurons in layers 2/3, along with the impaired spine morphology and density observed in both layer 2/3 and layer 5 of *Reln^del/+^* mice. These structural deficits contribute to mPFC dysfunction and may serve as a neurobiological basis for the social recognition impairments observed in these mice.

We observed a significant decrease in GABAergic interneurons in the mPFC of *Reln^del/+^* mice. Interestingly, although the number of Reelin-positive GABAergic neurons was decreased, the Reelin-negative GABAergic neuron number did not change. These findings imply that Reelin plays a vital role in maintaining the survival and function of specific GABAergic subtypes. Reelin is mainly produced by GABAergic interneurons and is essential for regulating cognitive function and neuronal plasticity [9], [10], [11]. This reduction of GABAergic interneurons may contribute to impaired inhibitory transmission in the PFC, a characteristic observed in SCZ [47].

PV-positive interneurons represent a major GABAergic subtype that exerts powerful perisomatic inhibition onto pyramidal neurons [48]. PV-positive interneurons are essential for the generation of gamma oscillations, which coordinate synchronous neuronal firing and support advanced functions such as cognitive abilities and social interaction [49]. Furthermore, we observed a reduction in PV boutons in the mPFC, indicating disrupted synaptic output from these interneurons. Since the VGLUT2/VGAT ratio on the soma of both pyramidal neurons and PV-positive interneurons remained unchanged (Fig. S2), E/I balance may be preserved at the individual cell level. Thus, the overall reduction in PV-positive interneurons and their synaptic boutons indicates weakened inhibitory output. This reduction, together with impaired structural integrity and synaptic organization of PV-positive interneurons, may compromise their inhibitory function and contribute to cortical dysfunction.

Previous studies demonstrated that mice treated with Reelin supplementation into the brain [11] or with overexpression of Reelin [50] exhibit enhanced cognition and long-term potentiation (LTP). The existing treatments for restoring the Reelin deficits involve injecting recombinant Reelin protein. Although recombinant Reelin protein has shown potential in preclinical models for restoring synaptic function and cognition, its rapid degradation within hours after injection poses a significant limitation [11], [51]. AAV vectors offer a compelling alternative for gene delivery to the central nervous system. These vectors facilitate efficient gene transfer, cross the blood-brain barrier, provide long-term expression, exhibit low immunogenicity, and allow targeted delivery to specific neuronal populations [52]. However, the large size of the Reelin sequence poses a challenge for insertion into AAV vectors. To circumvent this, a novel construct, R36, comprising Reelin repeats third and sixth, has been developed [7]. This construct effectively initiates intracellular signaling and enhances the Reelin pathway. A single intracerebroventricular administration of R36 protein into Reelin-deficient mice significantly improved cognitive function, providing preclinical evidence for its potential therapeutic application [7]. In Western blot analysis, no obvious myc band was detected in the mock group. This is mainly because the AAV-mock-Myc group express only the standalone myc tag (1.2 kDa), which is below the typical detection range of SDS-PAGE and Western blot. Furthermore, antibody binding efficiency to the isolated myc peptide is low, resulting in absence of a visible band. In contrast, the experimental group expresses the myc-tag targeting protein R36, with a molecular weight of 66 kDa, which is readily detected by the anti-myc antibody, producing a clear, specific band. Importantly, the successful rescue by AAV-R36-Myc injection of social novelty deficits in *Reln^del/+^* mice provides preclinical evidence for the functional role of Reelin in social cognition. These results demonstrate that restoring Reelin signaling in the mPFC can effectively improve social impairments, pointing to its potential role in SCZ treatment.

While our study advances understanding of the role of Reelin in mPFC function and SCZ-related social and cognitive behaviors, several limitations must be addressed. First, although our findings suggest that *Reln* deletion causes structural and functional deficits in the mPFC, the precise molecular mechanisms remain unclear. Future studies using electrophysiological recordings and transcriptomic analyses could provide deeper insights into how Reelin modulates synaptic plasticity and network activity. Second, while our rescue experiments demonstrate the therapeutic potential of R36, it remains unknown whether long-term Reelin restoration can fully reverse structural deficits. Such investigation will provide a more comprehensive understanding of how Reelin deficiency contributes to the pathophysiology in SCZ and may uncover additional therapeutic targets.

In summary, the *Reln^del/+^* mice exhibited abnormalities in the number and morphology of both excitatory and inhibitory neurons, including PV-positive interneurons in the mPFC. These changes were associated with deficits in social novelty behavior, mirroring some aspects of SCZ. The injection of AAV-R36-Myc into the mPFC of *Reln^del/+^* mice rescued social novelty deficits. Our findings suggest that *Reln^del/+^* mouse model of SCZ provides valuable insights into the neurobiological mechanisms underlying social cognitive impairments in this neurodevelopmental disorder.

## 5. CRediT authorship contribution statement

**Youyun Zhu**: Conceptualization, Methodology, Investigation, Formal analysis, Writing – original draft, Funding acquisition. **Kanako Kitagawa**: Data curation, Formal analysis, Investigation, Writing - review & editing. **Daisuke Mori**: Resources, Writing - review & editing. **Tetsuo Matsuzaki**: Resources, Writing - review & editing **Taku Nagai**: Writing - review & editing. **Toshitaka Nabeshima**: Writing - review & editing. **Sayaka Takemoto-Kimura**: Resources, Writing - review & editing. **Hiroaki Ikesue**: Writing - review & editing. **Norio Ozaki**: Funding acquisition, Resources, Writing - review & editing. **Hiroyuki Mizoguchi**: Supervision, Funding acquisition, Writing - review & editing. **Kiyofumi Yamada**: Conceptualization, Supervision, Funding acquisition, Project administration, Writing - review & editing.

## Acknowledgments

The authors wish to acknowledge the Division for Medical Research Engineering, Nagoya University Graduate School of Medicine, for the use of a BZ9000 bright-field microscopic (KEYENCE), two confocal laser scanning microscopes (TiE-A1R, Nikon; LSM880, Carl Zeiss AG), a cryostat (CM3050S; Leica), Imaris software (Oxford Instruments), and the staff of the Division of Experimental Animals, Nagoya University Graduate School of Medicine, for their technical support. The authors are grateful to Dr. Atsunobu Sagara and Dr. Masahito Sawahata for the kind help with animal experiment. Cartoons in Graphical abstract, Figures 1A and 5A were created with BioRender.com.

## References

[1] F. Tissir, A.M. Goffinet, Reelin and brain development, Nat. Rev. Neurosci. 4 (2003) 496–505. 10.1038/nrn1113.

[2] M. Ogawa, T. Miyata, K. Nakajimat, K. Yagyu, M. Seike, K. Ikenaka, H. Yamamoto, K. Mikoshibat, The reeler gene-associated antigen on cajal-retzius neurons is a crucial molecule for laminar organization of cortical neurons, Neuron 14 (1995) 899–912. 10.1016/0896-6273(95)90329-1.

[3] L.S. Turk, X. Kuang, V. Dal Pozzo, K. Patel, M. Chen, K. Huynh, M.J. Currie, D. Mitchell, R.C.J. Dobson, G. D’Arcangelo, W. Dai, D. Comoletti, The structure-function relationship of a signaling-competent, dimeric Reelin fragment, Structure 29 (2021) 1156–1170.e6. 10.1016/j.str.2021.05.012.

[4] M. Frotscher, Dual role of Cajal-Retzius cells and reelin in cortical development, Cell Tissue Res. 290 (1997) 315–322. 10.1007/s004410050936.

[5] T. Hiesberger, M. Trommsdorff, B.W. Howell, A. Goffinet, M.C. Mumby, J.A. Cooper, J. Herz, Direct Binding of Reelin to VLDL Receptor and ApoE Receptor 2 Induces Tyrosine Phosphorylation of Disabled-1 and Modulates Tau Phosphorylation, Neuron 24 (1999) 481–489. 10.1016/S0896-6273(00)80861-2.

[6] N.K. Morrill, Reelin Supplementation for the Rescue of Fragile X Syndrome., (n.d.).

[7] N.K. Morrill, Q. Li, A. Joly-Amado, E.J. Weeber, K.R. Nash, A novel Reelin construct, R36, recovered behavioural deficits in the heterozygous reeler mouse, Eur. J. Neurosci. 57 (2023) 1657–1670. 10.1111/ejn.15971.

[8] Y. Jossin, N. Ignatova, T. Hiesberger, J. Herz, C. Lambert De Rouvroit, A.M. Goffinet, The Central Fragment of Reelin, Generated by Proteolytic Processing *In Vivo*, Is Critical to Its Function during Cortical Plate Development, J. Neurosci. 24 (2004) 514–521. 10.1523/JNEUROSCI.3408-03.2004.

[9] S. Qiu, K. Korwek, A. Prattdavis, M. Peters, M. Bergman, E. Weeber, Cognitive disruption and altered hippocampus synaptic function in Reelin haploinsufficient mice, Neurobiol. Learn. Mem. 85 (2006) 228–242. 10.1016/j.nlm.2005.11.001.

[10] J. Herz, Y. Chen, Reelin, lipoprotein receptors and synaptic plasticity, Nat. Rev. Neurosci. 7 (2006) 850–859. 10.1038/nrn2009.

[11] J.T. Rogers, I. Rusiana, J. Trotter, L. Zhao, E. Donaldson, D.T.S. Pak, L.W. Babus, M. Peters, J.L. Banko, P. Chavis, G.W. Rebeck, H.-S. Hoe, E.J. Weeber, Reelin supplementation enhances cognitive ability, synaptic plasticity, and dendritic spine density, Learn. Mem. 18 (2011) 558–564. 10.1101/lm.2153511.

[12] A. Joly-Amado, N. Kulkarni, K.R. Nash, Reelin Signaling in Neurodevelopmental Disorders and Neurodegenerative Diseases, Brain Sci. 13 (2023) 1479. 10.3390/brainsci13101479.

[13] G.H. Lee, Z. Chhangawala, S. Von Daake, J.N. Savas, J.R. Yates, D. Comoletti, G. D’Arcangelo, Reelin Induces Erk1/2 Signaling in Cortical Neurons Through a Non-canonical Pathway, J. Biol. Chem. 289 (2014) 20307–20317. 10.1074/jbc.M114.576249.

[14] T.D. Folsom, S.H. Fatemi, The involvement of Reelin in neurodevelopmental disorders, Neuropharmacology 68 (2013) 122–135. 10.1016/j.neuropharm.2012.08.015.

[15] F. Impagnatiello, A.R. Guidotti, C. Pesold, Y. Dwivedi, H. Caruncho, M.G. Pisu, D.P. Uzunov, N.R. Smalheiser, J.M. Davis, G.N. Pandey, G.D. Pappas, P. Tueting, R.P. Sharma, E. Costa, A decrease of reelin expression as a putative vulnerability factor in schizophrenia, Proc. Natl. Acad. Sci. 95 (1998) 15718–15723. 10.1073/pnas.95.26.15718.

[16] A. Vílchez-Acosta, Y. Manso, A. Cárdenas, A. Elias-Tersa, M. Martínez-Losa, M. Pascual, M. Álvarez-Dolado, A.C. Nairn, V. Borrell, E. Soriano, Specific contribution of Reelin expressed by Cajal–Retzius cells or GABAergic interneurons to cortical lamination, Proc. Natl. Acad. Sci. 119 (2022) e2120079119. 10.1073/pnas.2120079119.

[17] D.A. Lewis, A.A. Curley, J.R. Glausier, D.W. Volk, Cortical parvalbumin interneurons and cognitive dysfunction in schizophrenia, Trends Neurosci. 35 (2012) 57–67. 10.1016/j.tins.2011.10.004.

[18] I. Kushima, B. Aleksic, M. Nakatochi, T. Shimamura, T. Shiino, A. Yoshimi, H. Kimura, Y. Takasaki, C. Wang, J. Xing, K. Ishizuka, T. Oya-Ito, Y. Nakamura, Y. Arioka, T. Maeda, M. Yamamoto, M. Yoshida, H. Noma, S. Hamada, M. Morikawa, Y. Uno, T. Okada, T. Iidaka, S. Iritani, T. Yamamoto, M. Miyashita, A. Kobori, M. Arai, M. Itokawa, M.-C. Cheng, Y.-A. Chuang, C.-H. Chen, M. Suzuki, T. Takahashi, R. Hashimoto, H. Yamamori, Y. Yasuda, Y. Watanabe, A. Nunokawa, T. Someya, M. Ikeda, T. Toyota, T. Yoshikawa, S. Numata, T. Ohmori, S. Kunimoto, D. Mori, N. Iwata, N. Ozaki, High-resolution copy number variation analysis of schizophrenia in Japan, Mol. Psychiatry 22 (2017) 430–440. 10.1038/mp.2016.88.

[19] M. Sawahata, D. Mori, Y. Arioka, H. Kubo, I. Kushima, K. Kitagawa, A. Sobue, E. Shishido, M. Sekiguchi, A. Kodama, R. Ikeda, B. Aleksic, H. Kimura, K. Ishizuka, T. Nagai, K. Kaibuchi, T. Nabeshima, K. Yamada, N. Ozaki, Generation and analysis of novel *Reln*ϑ deleted mouse model corresponding to exonic *Reln* deletion in schizophrenia, Psychiatry Clin. Neurosci. 74 (2020) 318–327. 10.1111/pcn.12993.

[20] Y. Tsuneura, M. Sawahata, N. Itoh, R. Miyajima, D. Mori, T. Kohno, M. Hattori, A. Sobue, T. Nagai, H. Mizoguchi, T. Nabeshima, N. Ozaki, K. Yamada, Analysis of Reelin signaling and neurodevelopmental trajectory in primary cultured cortical neurons with RELN deletion identified in schizophrenia, Neurochem. Int. 144 (2021) 104954. 10.1016/j.neuint.2020.104954.

[21] J. Liao, G. Dong, B. Wulaer, M. Sawahata, H. Mizoguchi, D. Mori, N. Ozaki, T. Nabeshima, T. Nagai, K. Yamada, Mice with exonic RELN deletion identified from a patient with schizophrenia have impaired visual discrimination learning and reversal learning in touchscreen operant tasks, Behav. Brain Res. 416 (2022) 113569. 10.1016/j.bbr.2021.113569.

[22] B. Rein, K. Ma, Z. Yan, A standardized social preference protocol for measuring social deficits in mouse models of autism, Nat. Protoc. 15 (2020) 3464–3477. 10.1038/s41596-020-0382-9.

[23] G. Paxinos, K.B.J. Franklin, Paxinos and Franklin’s The mouse brain in stereotaxic coordinates, Fifth edition, Academic Press, an imprint of Elsevier, London, 2019.

[24] K. Yamada, M. Takayanagi, H. Kamei, T. Nagai, M. Dohniwa, K. Kobayashi, S. Yoshida, T. Ohhara, K. Takauma, T. Nabeshima, Effects of memantine and donepezil on amyloid β-induced memory impairment in a delayed-matching to position task in rats, Behav. Brain Res. 162 (2005) 191–199. 10.1016/j.bbr.2005.02.036.

[25] F.W. Grillo, S. Song, L.M. Teles-Grilo Ruivo, L. Huang, G. Gao, G.W. Knott, B. Maco, V. Ferretti, D. Thompson, G.E. Little, V. De Paola, Increased axonal bouton dynamics in the aging mouse cortex, Proc. Natl. Acad. Sci. 110 (2013). 10.1073/pnas.1218731110.

[26] Y. Zhang, J.-T. Li, H. Wang, W.-P. Niu, C.-C. Zhang, Y. Zhang, X.-D. Wang, T.-M. Si, Y.-A. Su, Role of trace amine[associated receptor 1 in the medial prefrontal cortex in chronic social stress-induced cognitive deficits in mice, Pharmacol. Res. 167 (2021) 105571. 10.1016/j.phrs.2021.105571.

[27] H. Carceller, R. Guirado, E. Ripolles-Campos, V. Teruel-Marti, J. Nacher, Perineuronal Nets Regulate the Inhibitory Perisomatic Input onto Parvalbumin Interneurons and γ Activity in the Prefrontal Cortex, J. Neurosci. 40 (2020) 5008–5018. 10.1523/JNEUROSCI.0291-20.2020.

[28] K. Hada, B. Wulaer, T. Nagai, N. Itoh, M. Sawahata, A. Sobue, H. Mizoguchi, D. Mori, I. Kushima, T. Nabeshima, N. Ozaki, K. Yamada, Mice carrying a schizophrenia-associated mutation of the Arhgap10 gene are vulnerable to the effects of methamphetamine treatment on cognitive function: association with morphological abnormalities in striatal neurons, Mol. Brain 14 (2021) 21. 10.1186/s13041-021-00735-4.

[29] Y. Jossin, Reelin Functions, Mechanisms of Action and Signaling Pathways During Brain Development and Maturation, Biomolecules 10 (2020) 964. 10.3390/biom10060964.

[30] A.F.T. Arnsten, Prefrontal cortical network connections: key site of vulnerability in stress and schizophrenia, Int. J. Dev. Neurosci. 29 (2011) 215–223. 10.1016/j.ijdevneu.2011.02.006.

[31] M.F. Egan, D.R. Weinberger, Neurobiology of schizophrenia, Curr. Opin. Neurobiol. 7 (1997) 701–707. 10.1016/S0959-4388(97)80092-X.

[32] The Prefrontal Cortex, Elsevier, 2008. 10.1016/B978-0-12-373644-4.X0001-1.

[33] D.A. Lewis, G. González-Burgos, Neuroplasticity of Neocortical Circuits in Schizophrenia, Neuropsychopharmacology 33 (2008) 141–165. 10.1038/sj.npp.1301563.

[34] S. Wei, J. Jiang, D. Wang, J. Chang, L. Tian, X. Yang, X.-R. Ma, J.-W. Zhao, Y. Li, S. Chang, X. Chi, H. Li, N. Li, GPR158 in pyramidal neurons mediates social novelty behavior via modulating synaptic transmission in male mice, Cell Rep. 43 (2024) 114796. 10.1016/j.celrep.2024.114796.

[35] S.J. Kaar, I. Angelescu, T.R. Marques, O.D. Howes, Pre-frontal parvalbumin interneurons in schizophrenia: a meta-analysis of post-mortem studies, J. Neural Transm. 126 (2019) 1637–1651. 10.1007/s00702-019-02080-2.

[36] K. Kole, B.J.B. Voesenek, M.E. Brinia, N. Petersen, M.H.P. Kole, Parvalbumin basket cell myelination accumulates axonal mitochondria to internodes, Nat. Commun. 13 (2022) 7598. 10.1038/s41467-022-35350-x.

[37] W. Li, X. Guo, S. Xiao, Evaluating the relationship between reelin gene variants (rs7341475 and rs262355) and schizophrenia: A meta-analysis, Neurosci. Lett. 609 (2015) 42–47. 10.1016/j.neulet.2015.10.014.

[38] Z. Wang, Y. Hong, L. Zou, R. Zhong, B. Zhu, N. Shen, W. Chen, J. Lou, J. Ke, T. Zhang, W. Wang, X. Miao, Reelin gene variants and risk of autism spectrum disorders: An integrated meta[analysis, Am. J. Med. Genet. B Neuropsychiatr. Genet. 165 (2014) 192–200. 10.1002/ajmg.b.32222.

[39] A. Gee, P. Dazzan, A.A. Grace, G. Modinos, Corticolimbic circuitry as a druggable target in schizophrenia spectrum disorders: a narrative review, Transl. Psychiatry 15 (2025) 21. 10.1038/s41398-024-03221-2.

[40] J. Iafrati, M.J. Orejarena, O. Lassalle, L. Bouamrane, P. Chavis, Reelin, an extracellular matrix protein linked to early onset psychiatric diseases, drives postnatal development of the prefrontal cortex via GluN2B-NMDARs and the mTOR pathway, Mol. Psychiatry 19 (2014) 417–426. 10.1038/mp.2013.66.

[41] W.S. Liu, C. Pesold, M.A. Rodriguez, G. Carboni, J. Auta, P. Lacor, J. Larson, B.G. Condie, A. Guidotti, E. Costa, Down-regulation of dendritic spine and glutamic acid decarboxylase 67 expressions in the reelin haploinsufficient heterozygous reeler mouse, Proc. Natl. Acad. Sci. 98 (2001) 3477–3482. 10.1073/pnas.051614698.

[42] J. Leemhuis, E. Bouché, M. Frotscher, F. Henle, L. Hein, J. Herz, D.K. Meyer, M. Pichler, G. Roth, C. Schwan, H.H. Bock, Reelin Signals through Apolipoprotein E Receptor 2 and Cdc42 to Increase Growth Cone Motility and Filopodia Formation, J. Neurosci. 30 (2010) 14759–14772. 10.1523/JNEUROSCI.4036-10.2010.

[43] M. Matsuzaki, G.C.R. Ellis-Davies, T. Nemoto, Y. Miyashita, M. Iino, H. Kasai, Dendritic spine geometry is critical for AMPA receptor expression in hippocampal CA1 pyramidal neurons, Nat. Neurosci. 4 (2001) 1086–1092. 10.1038/nn736.

[44] R.R. Mahmmoud, S. Sase, Y.D. Aher, A. Sase, M. Gröger, M. Mokhtar, H. Höger, G. Lubec, Spatial and Working Memory Is Linked to Spine Density and Mushroom Spines, PLOS ONE 10 (2015) e0139739. 10.1371/journal.pone.0139739.

[45] V.E. Hammond, E. So, H.S. Cate, J.M. Britto, J.M. Gunnersen, S.-S. Tan, Cortical Layer Development and Orientation is Modulated by Relative Contributions of Reelin-Negative and-Positive Neurons in Mouse Chimeras, Cereb. Cortex 20 (2010) 2017–2026. 10.1093/cercor/bhp287.

[46] C.B. Klune, B. Jin, L.A. DeNardo, Linking mPFC circuit maturation to the developmental regulation of emotional memory and cognitive flexibility, eLife 10 (2021) e64567. 10.7554/eLife.64567.

[47] K. Ishii, K. Kubo, K. Nakajima, Reelin and Neuropsychiatric Disorders, Front. Cell. Neurosci. 10 (2016). 10.3389/fncel.2016.00229.

[48] H. Hu, J. Gan, P. Jonas, Fast-spiking, parvalbumin^+^ GABAergic interneurons: From cellular design to microcircuit function, Science 345 (2014) 1255263. 10.1126/science.1255263.

[49] B. Leitch, Parvalbumin Interneuron Dysfunction in Neurological Disorders: Focus on Epilepsy and Alzheimer’s Disease, Int. J. Mol. Sci. 25 (2024) 5549. 10.3390/ijms25105549.

[50] L. Pujadas, A. Gruart, C. Bosch, L. Delgado, C.M. Teixeira, D. Rossi, L. De Lecea, A. Martínez, J.M. Delgado-García, E. Soriano, Reelin Regulates Postnatal Neurogenesis and Enhances Spine Hypertrophy and Long-Term Potentiation, J. Neurosci. 30 (2010) 4636–4649. 10.1523/JNEUROSCI.5284-09.2010.

[51] D. Ibi, G. Nakasai, N. Koide, M. Sawahata, T. Kohno, R. Takaba, T. Nagai, M. Hattori, T. Nabeshima, K. Yamada, M. Hiramatsu, Reelin Supplementation Into the Hippocampus Rescues Abnormal Behavior in a Mouse Model of Neurodevelopmental Disorders, Front. Cell. Neurosci. 14 (2020) 285. 10.3389/fncel.2020.00285.

[52] D. Wang, P.W.L. Tai, G. Gao, Adeno-associated virus vector as a platform for gene therapy delivery, Nat. Rev. Drug Discov. 18 (2019) 358–378. 10.1038/s41573-019-0012-9.

