## Supplementary Data for "Cortical excitatory and inhibitory neuron deficits may underlie the cognitive and social impairments in a mouse model of schizophrenia with exonic *Reln* deletion"

**Supplementary Material**

**Fig. S1.** ***Reln*^del/+^ mice and *Reln*^del/del^ mice showed brain structure abnormality at development period.**

A. Representative images of body from 2-to 3-week-old wild type (WT), *Reln^del/+^* and *Reln^del/del^* mice
B. The body weight of WT, *Reln^del/+^* and *Reln^del/del^* mice (n =10 for WT mice; n = 10 for *Reln^del/+^* mice; n = 5 for *Reln^del/del^* mice)
C. Representative images of brain from 2-to 3-week-old WT, *Reln^del/+^* and *Reln^del/del^* mice
D. The brain weight of WT, *Reln^del/+^* and *Reln^del/del^* mice. (n =3 for WT mice; n = 3 for *Reln^del/+^* mice; n = 4 for *Reln^del/del^* mice)
E. Representative images of layer structure from the mPFC in WT, *Reln^del/+^* and *Reln^del/del^* mice. A multicolor confocal image taken from a mPFC region that was immunostained against Cux1 (Red, maker for layer 2/3) and Ctip2 (Green, marker for layer 5), indicating the clear boundaries of the layers. Scale bar, 200 μm.
F. The thickness of layer 2/3 and 5 in the medial prefrontal cortex (mPFC) from WT and *Reln^del/+^* mice. (n=11-13 slices from 3-4 mice per group)
G. Representative images of brain structure in WT, *Reln^del/+^* and *Reln^del/del^* mice. Scale bar, 500 μm.

For data represented as mean ± SEM; t tests. For all figures, ns. indicates not significant, *p < 0.05, ***p < 0.001, WT vs. mutant mice

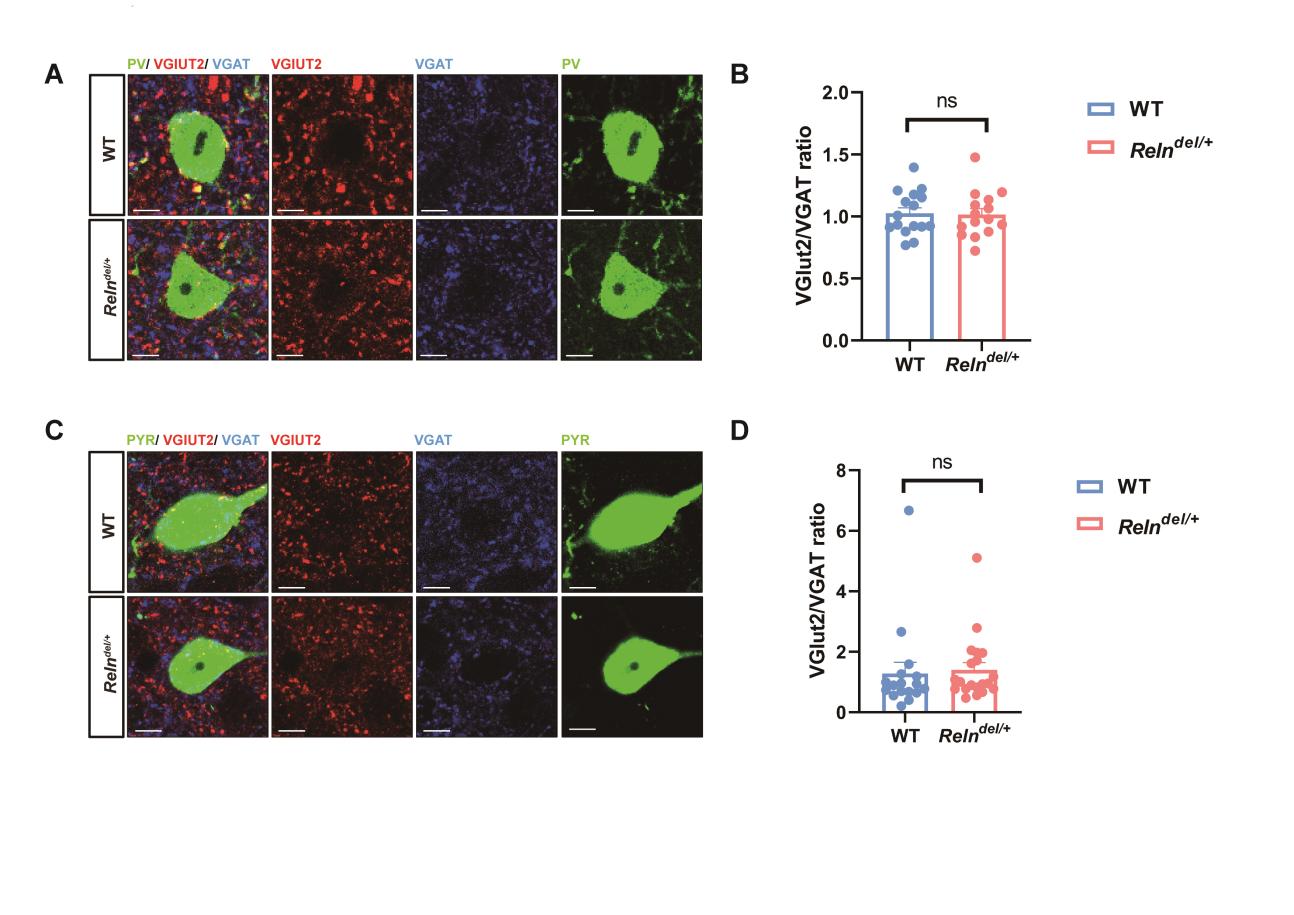

**Fig. S2. *Reln^del/+^* mice showed no change VGLUT2/VGAT ratio on the soma of PV-positive interneurons and pyramidal neurons**

A. Representative images of VGLUT2 and VGAT expression on the PV-positive interneuron’s soma of WT and *Reln^del/+^* mice (n=55-56 neurons from 4-5 mice per group). Scale bar, 5 μm.
B. The VGLUT2/VGAT ratio on the PV-positive interneuron’s soma of the medial prefrontal cortex (mPFC) from WT and *Reln^del/+^* mice.
C. Representative images of VGLUT2 and VGAT expression on the pyramidal neuron soma of WT and *Reln^del/+^* mice (n=75 neurons from 6 mice per group). Scale bar, 5 μm.
D. The VGLUT2/VGAT ratio on the pyramidal neuron soma of the mPFC from WT and *Reln^del/+^* mice.

For data represented as mean ± SEM; t tests. For all figures, ns. indicates not significant.

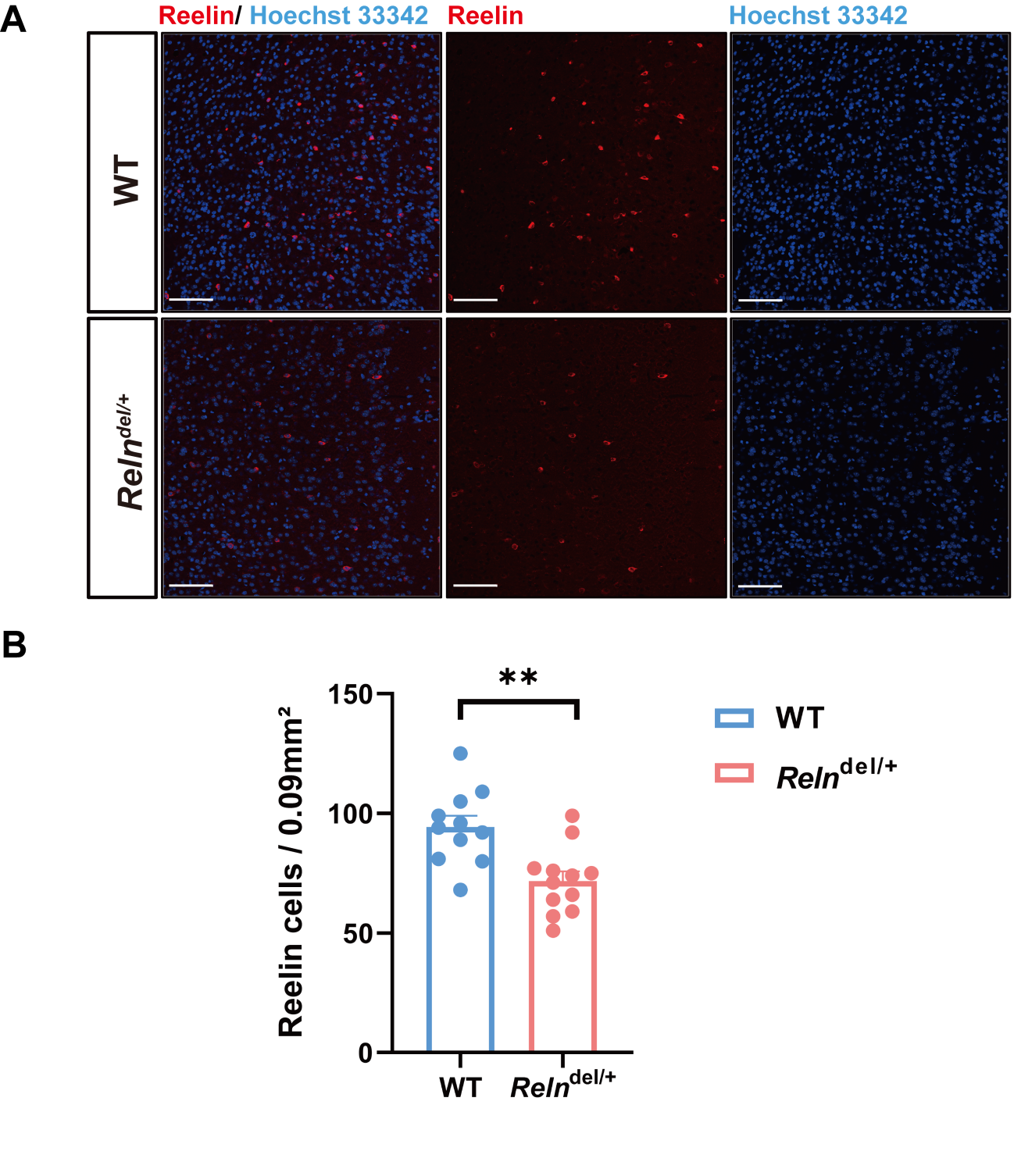

**Fig. S3. *Reln*^del/+^ mice showed that the number of Reelin-expression neurons in the mPFC was about 60% lower than in wild-type (WT) mice.**

A. Representative images of Reelin-expression neurons (Red) expression in the mPFC of WT and *Reln^del/+^* mice (n=11-12 slices from 4 mice per group). Scale bar, 100 μm.
B. The numbers of Reelin-expression neurons in the mPFC of WT and *Reln^del/+^* mice.

For data represented as mean ± SEM; t tests. For all figures, **p < 0.01, WT vs. mutant mice.

**Supplemental table 1 Antibody Information**

**Immunohistochemistry (IHC) - Primary Antibodies**

| Antibodies | Host Species | Clone/Code | Dilution | Company |
| --- | --- | --- | --- | --- |
| GABA | Rabbit | A2052-.2 ML | 1:1000 | Sigma-Aldrich |
| Parvalbumin (PV) | Rabbit | ab11427 | 1:500 | Abcam |
| CaMKIIα | Mouse | 05–532 | 1:1000 | Sigma-Aldrich |
| Reelin | Goat | AF3820 | 1:1000 | R&D Systems |
| Ctip2 | Rat | ab18465 | 1:1000 | Abcam |
| Cux1 | Rabbit | 11733-1-AP | 1:300 | Proteintech |
| VGAT | Mouse | #131011 | 1:1000 | Synaptic Systems |
| VGLUT2 | Guinea pig | #135404 | 1:1000 | Synaptic Systems |
| GFP | Rabbit | MBL598 | 1:500 | MBL |
| RFP | Rabbit | MBLPM005 | 1:500 | MBL |

**Immunohistochemistry (IHC) - Secondary Antibodies**

| Antibodies | Host Species | Code | Dilution | Company |
| --- | --- | --- | --- | --- |
| Donkey anti-mouse AF 594 | Mouse | A21203 | 1:1000 | Invitrogen |
| Donkey anti-goat AF 594 | Goat | A11058 | 1:1000 | Invitrogen |
| Donkey anti-mouse AF 647 | Mouse | A31571 | 1:1000 | Invitrogen |
| Goat anti-rabbit AF 488 | Rabbit | A11034 | 1:1000 | Invitrogen |
| Goat anti-guinea pig AF 568 | Guinea pig | A11075 | 1:1000 | Invitrogen |
| Rabbit anti-mouse AF 488 | Mouse | A21204 | 1:1000 | Invitrogen |
| Donkey anti-rabbit AF 488 | Rabbit | A21206 | 1:1000 | Invitrogen |
| Hoechst 33342 | — | 346–07951 | 1:2000 | Dojindo (Japan) |

**Western Blotting (WB) - Primary Antibodies**

| Antibodies | Host Species | Clone/Code | Dilution | Company |
| --- | --- | --- | --- | --- |
| Myc-Tag | Rabbit | 71D10 (#2278s) | 1:1000 | Cell Signaling |
| β-actin | Mouse | sc-47778 | 1:1000 | Santa Cruz |

**Western Blotting (WB) - Secondary Antibodies**

| Antibodies | Code | Dilution | Company |
| --- | --- | --- | --- |
| Rabbit IgG | 62–9520 | 1:10000 | Invitrogen |
| Mouse IgG | 5450–0011 | 1:10000 | Sera Care |

| **Supplemental table Statistics reporting by figure** | | | | | | |
| --- | --- | --- | --- | --- | --- | --- |
| **Figure** | **Number of samples** | | **Test used** | **Degree of freedom and F/t/P value** | **Post-hoc test** | **Significance** |
| **Figure 1.C** | Female: Control=6 | | Two-way ANOVA | Interaction: F (1, 44) = 3.530, P=0.0669 | Tukey's multiple comparisons test | WT: M1 vs.O2: P=<0.0001 |
|  | Female: Reln^del/+^=6 | |  | Social stimulation: F (1, 44) = 108.6. P<0.0001 |  |  |
|  | Male: Control=6 | |  | Gene: F (1, 44) = 2.004, P=0.1640 |  | Reln^del/+^: M1 vs.O2: P=<0.0001 |
|  | Male: Reln^del/+^=6 | |  |  |  |  |
| **Figure 1.D** | Female: Control=6 | | Unpaired t test |  |  | WT vs. Reln^del/+^: P=0.2665 |
|  | Female: Reln^del/+^=6 | |  |  |  |  |
|  | Male: Control=6 | |  |  |  |  |
|  | Male: Reln^del/+^=6 | |  |  |  |  |
| **Figure 1.G** | Female:Control=6 | | Two-way ANOVA | Interaction: F (1, 44) = 15.41, P=0.0003 | Tukey's multiple comparisons test | WT:M1 vs.M2: P=<0.0001 |
|  | Female: Reln^del/+^=6 | |  | Social stimulation: F (1, 44) = 24.37, P<0.0001 |  |  |
|  | Male: Control=6 | |  | Gene: F (1, 44) = 0.4846, P=0.4900 |  | Reln^del/+^:M1 vs.M2: P=0.8908 |
|  | Male: Reln^del/+^=6 | |  |  |  |  |
| **Figure 1.H** | Female: Control=6 | | Unpaired t test |  |  | Contrl vs. Reln^del/+^: P=0.0242 |
|  | Female: Reln^del/+^=6 | |  |  |  |  |
|  | Male: Control=6 | |  |  |  |  |
|  | Male: Reln^del/+^=6 | |  |  |  |  |
| **Figure 2.B** | 22 neurons from 5 mice per group | | Unpaired t test |  |  | WT vs. Reln^del/+^: P=<0.0001 |
| **Figure 2.C** | 22 neurons from 5 mice per group | | Unpaired t test |  |  | Mushroom type: WT vs. Reln^del/+^: P=0.0069 |
|  |  |  |  |  |  | Stuby type: WT vs. Reln^del/+^: P=0.0483 |
|  |  |  |  |  |  | Long thin type: WT vs. Reln^del/+^: P=0.0501 |
|  |  |  |  |  |  | Flipodia type: WT vs. Reln^del/+^: P=0.0008 |
| **Figure 2.E** | 23-25 neurons from 5 mice per group | | Unpaired t test |  |  | WT vs. Reln^del/+^: P=<0.0001 |
| **Figure 2.F** | 23-25 neurons from 5 mice per group | | Unpaired t test |  |  | Mushroom type: WT vs. Reln^del/+^: P=0.0011 |
|  |  |  |  |  |  | Stuby type: WT vs. Reln^del/+^:P=0.0238 |
|  |  |  |  |  |  | Long thin type: WT vs. Reln^del/+^: P=0.0511 |
|  |  |  |  |  |  | Flipodia type: WT vs. Reln^del/+^: P=0.0082 |
| **Figure 3.D** | 9 slices from 3 mice per group | | Unpaired t test |  |  | CaMKII-positive pyramidal neurons in Layer 2/3: WT vs. Reln^del/+^: P=0.0072 |
|  |  |  |  |  |  | CaMKII-positive pyramidal neurons in Deep Layer 2/3: WT vs. Reln^del/+^: P=0.0028 |
|  |  |  |  |  |  | CaMKII-positive pyramidal neurons in Layer 5: WT vs. Reln^del/+^: P>0.9999 |
| **Figure 4.B** | 9 slices from 3 mice per group | | Unpaired t test |  |  | GABAergic interneurons: WT vs. Reln^del/+^: P=0.0054 |
| **Figure 4.C** | 9 slices from 3 mice per group | | Unpaired t test |  |  | Reelin-GABAergic neurons WT vs. Reln^del/+^: P=0.0224 |
| **Figure 4.D** | 9 slices from 3 mice per group | | Unpaired t test |  |  | Reelin negative-GABAergic neurons WT vs. Reln^del/+^:P=0.3565 |
| **Figure 5.C** | 29-31 neurons from 5 mice per group | | Unpaired t test |  |  | PV- positive interneurons: WT vs. Reln^del/+^: P=<0.0001 |
| **Figure 5.E** | 29-31 neurons from 5 mice per group | | Unpaired t test |  |  | Total dendritic length: WT vs. Reln^del/+^: P=<0.0001 |
| **Figure 5.F** | Control: 18 neurons from 5 mice | | Two-way ANOVA | Interaction: F (25, 1225) = 4.665, P<0.0001 | Bonferroni's multiple comparisons test | Dendritic branches of PV-positive interneurons: 10μm WT vs. Reln^del/+^: P=<0.0001 |
|  |  |  |  | Distance: F (25, 1225) = 228.0, P<0.0001 |  | Dendritic branches of PV-positive interneurons: 20μm WT vs. Reln^del/+^: P=<0.0001 |
|  | Reln^del/+^：31 neurons from 5 mice | |  | Gene: F (1, 49) = 4.388, P=0.0414 |  | Dendritic branchesof of PV-positive interneurons: 40μm WT vs. Reln^del/+^: P=0.0056 |
|  |  |  |  | Subject factor: F (49, 1225) = 11.02, P<0.0001 |  | Dendritic branches of PV-positive interneurons: 70μm WT vs. Reln^del/+^: P=0.0416 |
| **Figure5.G** | Control: 18 neurons from 5 mice | | Two-way ANOVA | Interaction: F (27, 1296) = 5.147, P<0.0001 | Bonferroni's multiple comparisons test | Dendritic intersections of PV-positive interneurons: 10μm WT vs. Reln^del/+^: P=<0.0001 |
|  |  |  |  | Distance: F (27, 1296) = 228.5, P<0.0001 |  | Dendritic intersections of PV-positive interneurons: 20μm WT vs. Reln^del/+^: P=<0.0001 |
|  | Reln^del/+^：31 neurons from 5 mice | |  | Gene: F (1, 48) = 4.779, P=0.0337 |  | Dendritic intersections of PV-positive interneurons: 40μm WT vs. Reln^del/+^: P=0.0019 |
|  |  |  |  | Subject factor: F (48, 1296) = 7.716 P<0.0001 |  | Dendritic intersections of PV-positive interneurons: 60μm WT vs. Reln^del/+^: P=0.0138 |
| **Figure 5.I** | 23-30 axons from 5 mice per group | | Unpaired t test |  |  | PV boutons: WT; PV-Cre vs. Reln^del/+^; PV-Cre: P=0.0028 |
| **Figure 6.D** | WT mock | Female=5 | Two-way ANOVA | Interaction: F (3, 78) = 0.2522, P=0.8595 | Sidak's multiple comparisons test | WT mock: M1 vs.O2: P= 0.0004 |
|  |  | Male=6 |  | Social stimulation: F (1, 78) = 45.02, P<0.0001 |  |  |
|  | WT AAV-R36-Myc | Female=5 |  | Gene: F (3, 78) = 0.8045, P=0.4951 |  | WT AAV-R36-Myc: M1 vs.O2: P=0.0029 |
|  |  | Male=5 |  |  |  |  |
|  | Reln^del/+^ mock | Female=6 |  |  |  | Reln^del/+^ mock: M1 vs.O2: P=0.0082 |
|  |  | Male=6 |  |  |  |  |
|  | Reln^del/+^ AAV-R36-Myc | Female=5 |  |  |  | Reln^del/+^ AAV-R36-Myc: M1 vs.O2: P=<0.0001 |
|  |  | Male=5 |  |  |  |  |
| **Figure 6.E** | WT mock | Female=5 | Ordinary one-way ANOVA | Treatment (between columns): F (3, 39) = 0.6612, P=0.5809 | Sidak's multiple comparisons test | WT mock vs WT AAV-R36-Myc: P=0.3856 |
|  |  | Male=6 |  |  |  |  |
|  | WT AAV-R36-Myc | Female=5 |  |  |  |  |
|  |  | Male=5 |  |  |  |  |
|  | Reln^del/+^ mock | Female=6 |  |  |  | Reln^del/+^ mock vs Reln^del/+^ AAV-R36-Myc: P=0.7845 |
|  |  | Male=6 |  |  |  |  |
|  | Reln^del/+^ AAV-R36-Myc | Female=5 |  |  |  |  |
|  |  | Male=5 |  |  |  |  |
| **Figure 6.F** | WT mock | Female=5 | Two-way ANOVA | Interaction: F (3, 78) = 2.221, P=0.0923 | Sidak's multiple comparisons test | WT mock: M1 vs.O2: P= 0.0072 |
|  |  | Male=6 |  | Social stimulation: F (1, 78) = 34.63,P<0.0001 |  |  |
|  | WT AAV-R36-Myc | Female=5 |  | Gene: F (3, 78) = 1.775, P=0.1588 |  | WT AAV-R36-Myc: M1 vs.O2: P=0.0004 |
|  |  | Male=5 |  |  |  |  |
|  | Reln^del/+^ mock | Female=6 |  |  |  | Reln^del/+^ mock: M1 vs.O2: P>0.9999 |
|  |  | Male=6 |  |  |  |  |
|  | Reln^del/+^ AAV-R36-Myc | Female=5 |  |  |  | Reln^del/+^ AAV-R36-Myc: M1 vs.O2: P=0.0010 |
|  |  | Male=5 |  |  |  |  |
| **Figure 6.G** | WT mock | Female=5 | Ordinary one-way ANOVA | Treatment (between columns): F (3, 39) = 2.194, P=0.1042 | Sidak's multiple comparisons test | WT mock vs WT AAV-R36-Myc: P=0.9998 |
|  |  | Male=6 |  |  |  |  |
|  | WT AAV-R36-Myc | Female=5 |  |  |  |  |
|  |  | Male=5 |  |  |  |  |
|  | Reln^del/+^ mock | Female=6 |  |  |  | Reln^del/+^ mock vs Reln^del/+^ AAV-R36-Myc: P=0.0411 |
|  |  | Male=6 |  |  |  |  |
|  | Reln^del/+^ AAV-R36-Myc | Female=5 |  |  |  |  |
|  |  | Male=5 |  |  |  |  |
| **Figure S1.B** | Control=10 | | Ordinary one-way ANOVA | Treatment (between columns): F (2, 22) = 14.32, P=0.0001 | Sidak's multiple comparisons test | Body weight: WT vs. *Reln^del/+^*: P=0.9446 |
|  | *Reln^del/+^*=10 | |  |  |  | Body weight: WT vs. *Reln^del/del^*: P=0.0001 |
|  | *Reln^del/del^*=5 | |  |  |  |  |
| **Figure S1.D** | Control=3 | | Ordinary one-way ANOVA | Treatment (between columns): F (2, 7) = 5.742, P=0.0334 | Sidak's multiple comparisons test | Brain weight: WT vs. *Reln^del/+^*: P=0.9633 |
|  | *Reln^del/+^*=3 | |  |  |  | Brain weight: WT vs. *Reln^del/del^*: P=0.0400 |
|  | *Reln^del/del^*=4 | |  |  |  |  |
| **Figure S1.F** | 11-14 neurons from 3 mice per group | | Unpaired t test |  |  | Layer 2/3: WT vs. *Reln^del/+^*: P=0.0436 |
| **Figure S2.B** | 55 -56 neurons from 4-5 mice per group | | Unpaired t test |  |  | Layer 2/3: WT vs. *Reln^del/+^*: P=0.7195 |
|  |  |  |  |  |  | VGLUT2/VGAT ratio on the soma of Pv- positive interneurons: WT vs. *Reln^del/+^*: P=0.8731 |
| **Figure S2.D** | 75 neurons from 6 mice per group | | Unpaired t test |  |  | VGLUT2/VGAT ratio on the soma of pyrimidal neurons: WT vs. *Reln^del/+^*: P=0.7789 |
| **Figure S3.B** | 11-12 slices neurons from 3 mice per group | | Unpaired t test |  |  | Reelin- expression cells: WT vs. *Reln^del/+^*: P=0.0014 |
